# CloneVI: A deep generative model for deciphering the relationships between cell state and mitochondrial genotype

**DOI:** 10.64898/2026.11.03.764043

**Authors:** Taichi Hibi, Yasuhiro Kojima, Shuto Hayashi, Haruka Hirose, Chikara Mizukoshi, Teppei Shimamura

## Abstract

Cellular diversity arises from both genetic lineages and non-genetic regulation of cell states, and knowing which cell states are tied to lineages is central to understanding how clones shape disease. Single-cell multi-omics technologies measure mitochondrial DNA mutations together with gene expression or chromatin accessibility in the same cells, but clones are usually inferred from mutations alone, making such clones difficult to identify. We developed CloneVI, a deep generative model that infers clonal structure from a latent cell state derived from gene expression or chromatin accessibility. It estimates the clonal probability of each cell from this cell state, together with the allele frequency of each clone by reconstructing the observed mutation counts. In simulated data, CloneVI recovered the ground-truth clonal structure in both single-clone and mixed-clone cell populations. In clonal hematopoiesis single-cell RNA-seq data with enriched mitochondrial transcripts, the inferred clones were associated with specific mitochondrial DNA mutations, and some of them showed high expression of a T-cell receptor variable-region gene. In chronic lymphocytic leukemia mitochondrial single-cell ATAC-seq data, CloneVI separated populations with different heteroplasmy levels and identified increased PAX9 accessibility. In a multi-donor ovarian tumor cohort, CloneVI captured mutant-cell populations enriched in particular cell states, and a differential test weighted by the clonal probability revealed the transcription factor activity of the cell state in which the mutant cells concentrated. CloneVI thus identifies populations defined by mitochondrial DNA mutations and associated with cell states in single-cell RNA-seq and ATAC-seq data. CloneVI is available at https://github.com/TaichiHIBI/cloneVI.

## Introduction

Intratumoral heterogeneity is an important factor contributing to the complexity of cancer(1). Through the accumulation of genetic mutations, cancer cells form lineages with distinct genomes, commonly referred to as clones(2). These genetic differences contribute to variation in cellular behavior, including the acquisition of drug resistance and metastatic potential(1, 3). Understanding the composition and properties of individual clones is therefore important for understanding cancer evolution.

Considerable progress has been made in describing genetic heterogeneity. Whole-exome sequencing and single-cell DNA sequencing can reveal the clonal composition of tumors from somatic mutations(4, 5). At the same time, non-genetic heterogeneity also contributes to tumor diversity(6– 8). Single-cell RNA-seq (scRNA-seq) and single-cell ATAC-seq (scATAC-seq) have shown that cells with the same genome can assume diverse transcriptional and chromatin states. Cellular diversity therefore reflects both genetic lineage and non-genetic cell state. Because cells of the same lineage need not share a state, knowing which cell states are tied to lineages is central to understanding the properties of clones, and relating the two requires measuring them in the same cells.

Recent single-cell multi-omics technologies make it possible to measure genetic mutations from scRNA-seq or scATAC-seq data together with gene expression or chromatin accessibility in the same cells(9, 10). Mitochondrial DNA (mtDNA) mutations can be measured in the same way and used as endogenous clonal markers(11–15). mtDNA has a high copy number in each cell, making mtDNA mutations detectable in single-cell assays even when nuclear somatic variants are sparsely covered(16). Statistical methods have therefore been developed to identify informative mtDNA variants and infer clonal structure from single-cell data(11, 12, 17, 18).

Building on such genetic information, Cardelino and Clone-Tracer associate genetically defined clones with transcriptional states(19, 20). These methods first construct clonal structure from genotype information and then map gene-expression profiles to the resulting clones. Consequently, the clones are defined independently of cell states, and when the cells of a clone are in different states, the states that characterize the clone may be obscured.

Here, we propose CloneVI, a deep generative model that integrates genetic and non-genetic heterogeneity. CloneVI first infers cell states from gene expression or chromatin accessibility. It then estimates the probability of each clone given the cell state, together with the mtDNA allele frequencies that characterize each clone.

We first tested whether CloneVI recovers the expected clonal structure in simulated data. We then applied it to two small datasets of different modalities, mitochondrial alteration enrichment from single-cell transcriptomes to establish relatedness (MAESTER)(12) data from clonal hematopoiesis and mitochondrial single-cell ATAC-seq (mtscATAC-seq)(11) data from chronic lymphocytic leukemia (CLL), and finally to a large mtscATAC-seq cohort of ovarian tumors from five donors. CloneVI captures mutant-cell populations (cells that carry a given mtDNA variant) associated with particular cell states, and characterizes those cell states.

## Results

### CloneVI integrates cell state and mtDNA mutation information

CloneVI is a variational autoencoder (VAE)(21) that combines genetic and non-genetic information (Fig. 1; Supplementary Fig. S1). An encoder compresses the gene expression or chromatin accessibility counts **x**_*c*_ of each cell into a low-dimensional latent variable **z**_*c*_, and a decoder reconstructs the input counts **x**_*c*_ from **z**_*c*_. We refer to **z**_*c*_ as the cell state.

**Fig. 1.**
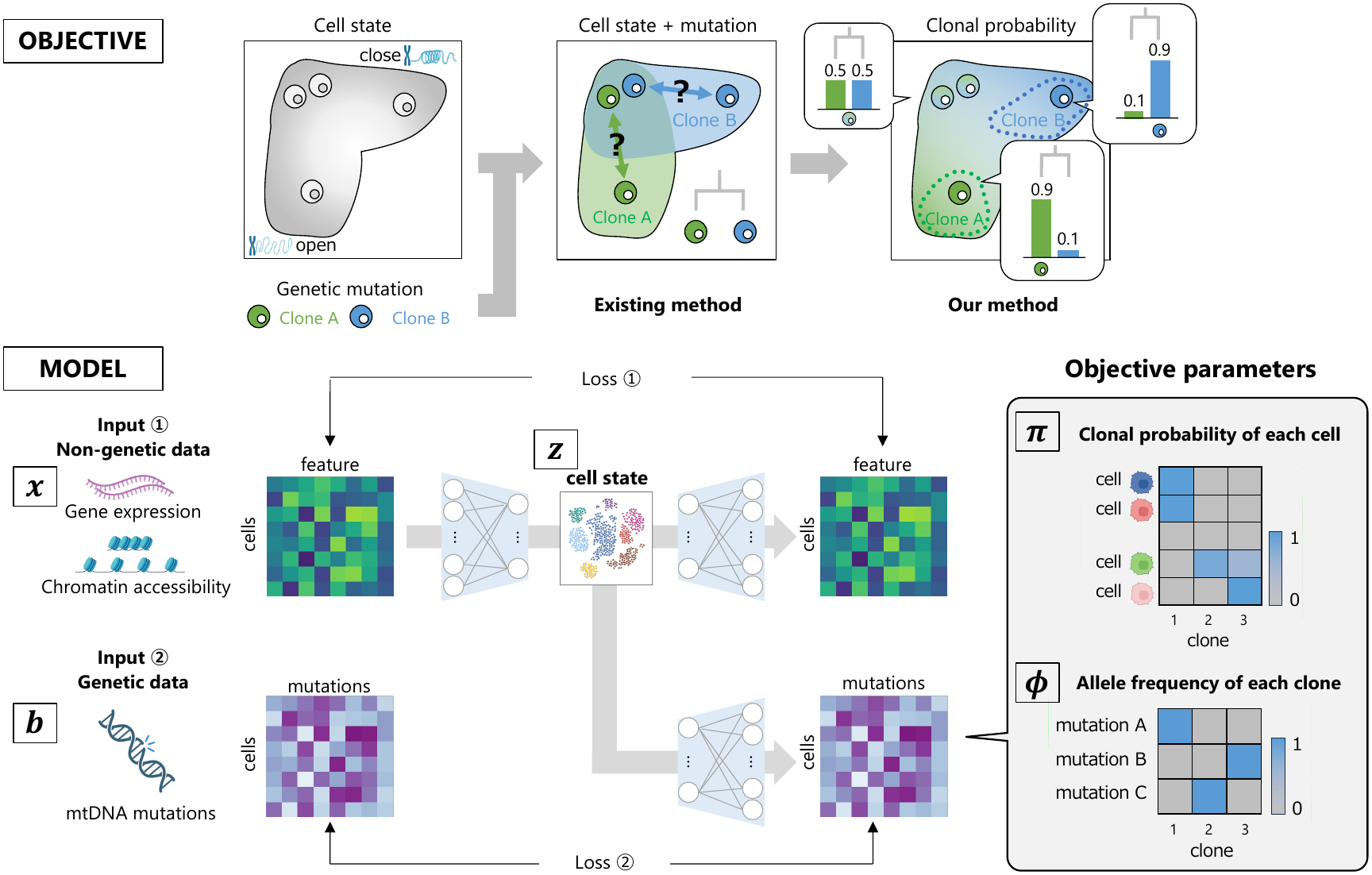
Overview of CloneVI. Top, the objective: existing methods assign cells to clones from their mutations, whereas CloneVI gives each cell a probability of belonging to each clone within the cell-state space. Bottom, the model: gene expression or chromatin accessibility **x** and mtDNA mutation counts **b** are provided for the same cells. CloneVI first encodes the non-genetic data into a latent cell state **z**. It then estimates the clonal probability ***π*** of each cell and the clone-specific allele frequency ***ϕ*** of each clone by reconstructing the observed mtDNA mutation counts. Icons: Created in BioRender. Hibi, T. (2026) https://BioRender.com/53seir5. The icons are not covered by the terms of the Creative Commons license of this publication; for permission to reuse, please contact the rights holder.

From the cell state, CloneVI then reconstructs the mtDNA mutation counts **b**_*c*_ of each cell. It estimates the clonal probability ***π***_*c*_ of each cell and the clone-specific allele frequency ***ϕ***_*k*_ of each clone *k* together, by maximizing the likelihood of the observed counts. Because ***π***_*c*_ is computed from the cell state alone, without the mutation counts of the cell itself, it gives the probability of each clone given the cell state, and the clonal probabilities form a landscape over the cell-state space. Throughout, a clone denotes one of these *K* inferred components; we call a lineage of cells sharing mutations a genetic clone, and the cells observed to carry a given variant a mutant-cell population.

Conventional methods define genetic clones from mutations and then treat gene expression or chromatin accessibility as properties of those clones. In contrast, CloneVI estimates clonal probabilities from the cell state.

CloneVI has four principal features:

1. **Continuous and probabilistic clone inference**: Clonal probabilities change continuously across the cell-state space, so cells near an ambiguous clone boundary, or in a cell state shared by several clones, are not forced into one clone.
2. **Selective capture of cell-state-enriched populations**: Mutant-cell populations that are enriched in particular cell states are preferentially captured as clones.
3. **Separation of clone-specific and background mutations**: The allele frequency of each clone can be split into a component shared by all clones and a clone-specific component (Methods).
4. **Separation by heteroplasmy**: Because ***ϕ*** is an allele frequency rather than a binary mutation status,

CloneVI can separate clones that carry the same mutation at different heteroplasmy levels.

The same model applies whether the cell-state input is gene expression or chromatin accessibility, so it can be used for both scRNA-seq and scATAC-seq data paired with mtDNA mutations.

### CloneVI recovers clonal probabilities in simulated data

We first used simulated data to test whether CloneVI recovers the expected clonal structure. The simulation was based on the Scanpy PBMC3k scRNA-seq dataset. For each expression cluster, we specified the probability that its cells belonged to Clone 0 or Clone 1 (Fig. 2a). Some clusters contained cells of only one clone, and others contained cells of both.

**Fig. 2.**
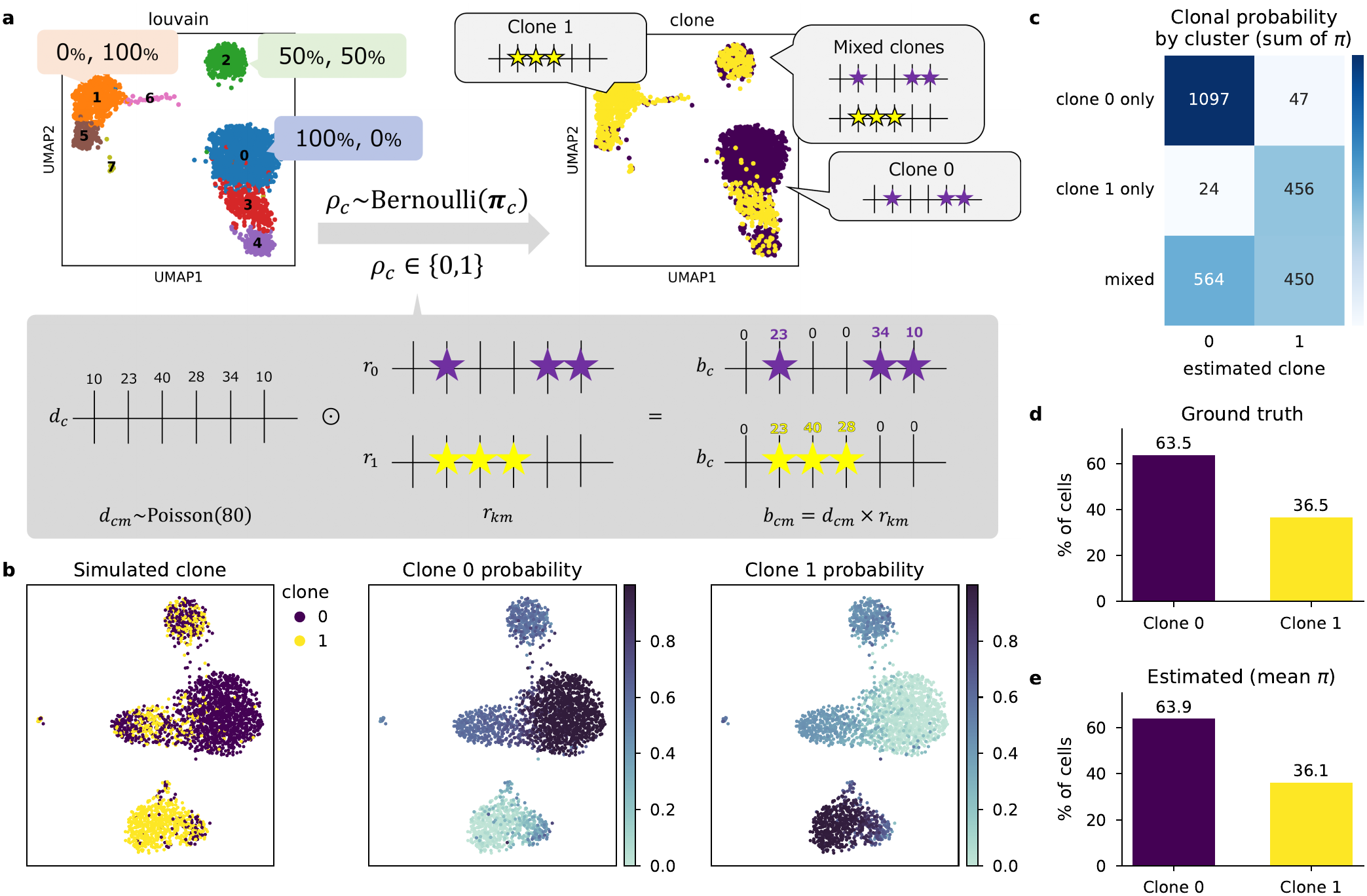
Validation using simulated data. **a**, Cells in the PBMC3k dataset were assigned to one of two clones according to probabilities defined for each expression cluster, and clone-specific mutation profiles were generated. **b**, Ground-truth clone labels and clonal probabilities estimated by CloneVI. **c**, Summed estimated clonal probabilities in the clusters containing only Clone 0, only Clone 1, or both (mixed). **d**,**e**, Ground-truth and estimated clone proportions in the complete dataset.

Each cell was assigned to a clone according to the probability of its cluster and given the mtDNA mutation profile of that clone (Supplementary Fig. S2). CloneVI was fitted to the expression counts and the mutation counts, without the clone labels.

Cells in clusters containing only one clone received high probability for that clone, whereas in mixed clusters the probabilities were split between the two clones (Fig. 2b,c). The mean clonal probabilities also matched the overall fraction of cells in each clone (Fig. 2d,e). These results indicate that CloneVI recovers both the dominant clone of each cluster and the degree of mixing as continuous probabilities.

### CloneVI identifies mutation-associated cell states in MAESTER data

We next applied CloneVI to MAESTER data, in which gene expression and mtDNA mutations were measured in the same cells of a patient with clonal hematopoiesis, using the 26 mtDNA variants analyzed in the original study(12).

CloneVI inferred six clones (Clones 0–5; Supplementary Fig. S3b). Clones 1–5 each carried particular mtDNA variants (Fig. 3b) and had their highest probability where the mutant cells of those variants were observed (Fig. 3c; Supplementary Fig. S3b,d).

**Fig. 3.**
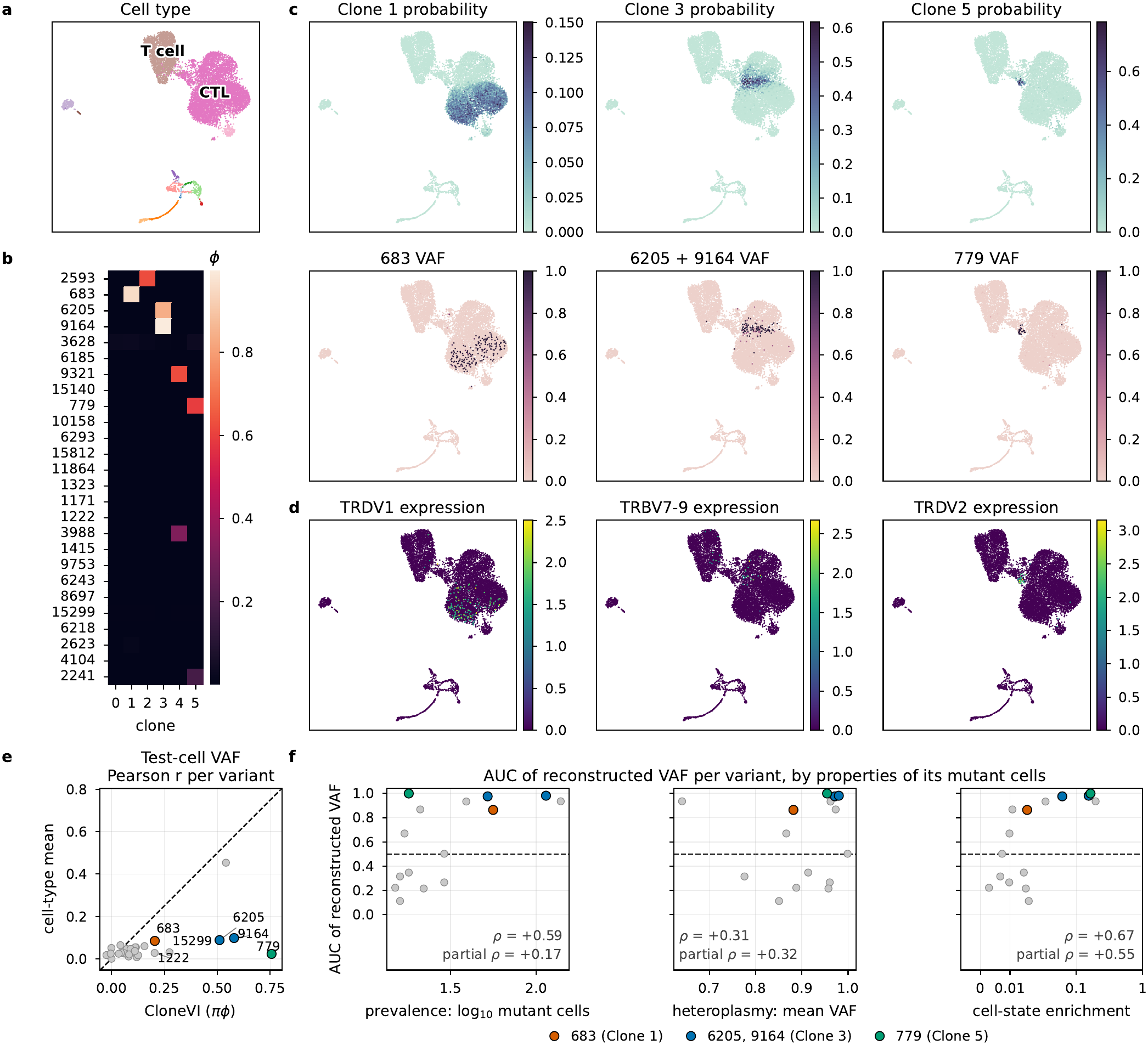
CloneVI applied to MAESTER data, **a**, Cell types on the UMAP of the cell-state space; T cells and cytotoxic T lymphocytes (CTL) are labeled, and all cell types are shown in Supplementary Fig. S3a. **b**, Clone-specific allele frequencies *ϕ* of the inferred clones, **c**, Clonal probabilities of Clones 1,3 and 5 (top) and observed variant allele frequency (VAF) of the variants associated with each clone (bottom), **d**, Expression of the T-cell receptor (TCR) variable-region gene selected for each clone, **e**, Pearson correlation between observed and reconstructed VAF in test cells for each variant, comparing reconstruction by CloneVI (*πϕ*) with the mean VAF of each cell type in training and validation cells, **f**, AUG with which the reconstructed VAF ranks mutant above non-mutant cells for the 15 variants, against prevalence, mean VAF of mutant cells and cell-state enrichment; *ρ*, Spearman correlation; partial *ρ*, partial Spearman correlation adjusted for the other two properties; colored points, variants of Clones 1,3 and 5.

Clones 3 and 5 each showed high expression of a particular T-cell receptor (TCR) variable-region gene, TRBV7-9 and TRDV2, in the same region of the cell-state space as their high clonal probability (Fig. 3c,d; Supplementary Fig. S3c). The probability of Clone 1 was instead spread thinly over the cytotoxic T-cell region (at most 0.15; Fig. 3a,c), where the mutant cells of its variant 683 were scattered.

We next asked whether the inferred clones could predict the observed variant allele frequency (VAF) of test cells. For each cell and variant, we computed the reconstructed VAF ***πϕ***, the VAF expected from the clonal probabilities of the cell and the allele frequencies of the clones (Methods). As a baseline, each test cell was given the mean VAF of its cell type in the training and validation cells. The reconstructed VAF agreed with the observed VAF better than the baseline for 21 of the 26 variants (mean Pearson r = 0.17 versus 0.05; Fig. 3e).

Finally, we examined which mutant-cell populations are captured by CloneVI. We regarded a population as captured when its mutant cells receive a higher reconstructed VAF than the other cells, and measured this for each variant by the area under the receiver operating characteristic curve (AUC): the probability that a mutant cell has a higher reconstructed VAF than a non-mutant cell (0.5, no separation; 1, complete separation). We compared the AUC with three properties of the variant: prevalence, mean VAF of mutant cells, and cell-state enrichment, which measures how much more often mutant cells neighbor other mutant cells in the cell-state space than expected at random (Methods). Across the 15 variants with enough mutant and non-mutant cells (Methods), the AUC was positively correlated with cell-state enrichment (Spearman correlation *ρ* = +0.67; Fig. 3f). Prevalence was also correlated with the AUC (*ρ* = +0.59) but was itself correlated with enrichment (*ρ* = +0.60). After adjustment for the other two properties, enrichment kept the largest partial Spearman correlation (partial *ρ* = +0.55), and those of prevalence an mean VAF were smaller but positive (partial p = +0.17 and +0.32). The variants of Clones 1,3 and 5 had AUCs of 0.86-1.00, whereas the five variants with almost no enrichment (below 0.01) had AUCs of 0.22-0.67. These results suggest that CloneVI preferentially captures mutant-cell populations that are enriched in particular cell states; with 15 variants, however, the three properties could not be fully separated, and we return to this question in a larger cohort below.

Together, these results indicate that the clones inferred by CloneVI from the MAESTER data link particular mtDNA variants to the cell states of their mutant cells, and that the clonal probabilities, estimated from the cell state alone, predict the VAF of unseen cells better than the mean VAF of their cell type.

### CloneVI identifies clone-associated chromatin states in mtscATAC-seq data

To test whether CloneVI could also identify clones associated with chromatin states, we analyzed mtscATAC-seq data from a patient with CLL(11). From 18 mtDNA variants, CloneVI inferred nine clones, with their clonal probabilities across the cell-state space and their allele frequencies (Fig. 4a-c; Supplementary Fig. S4a-c).

**Fig. 4.**
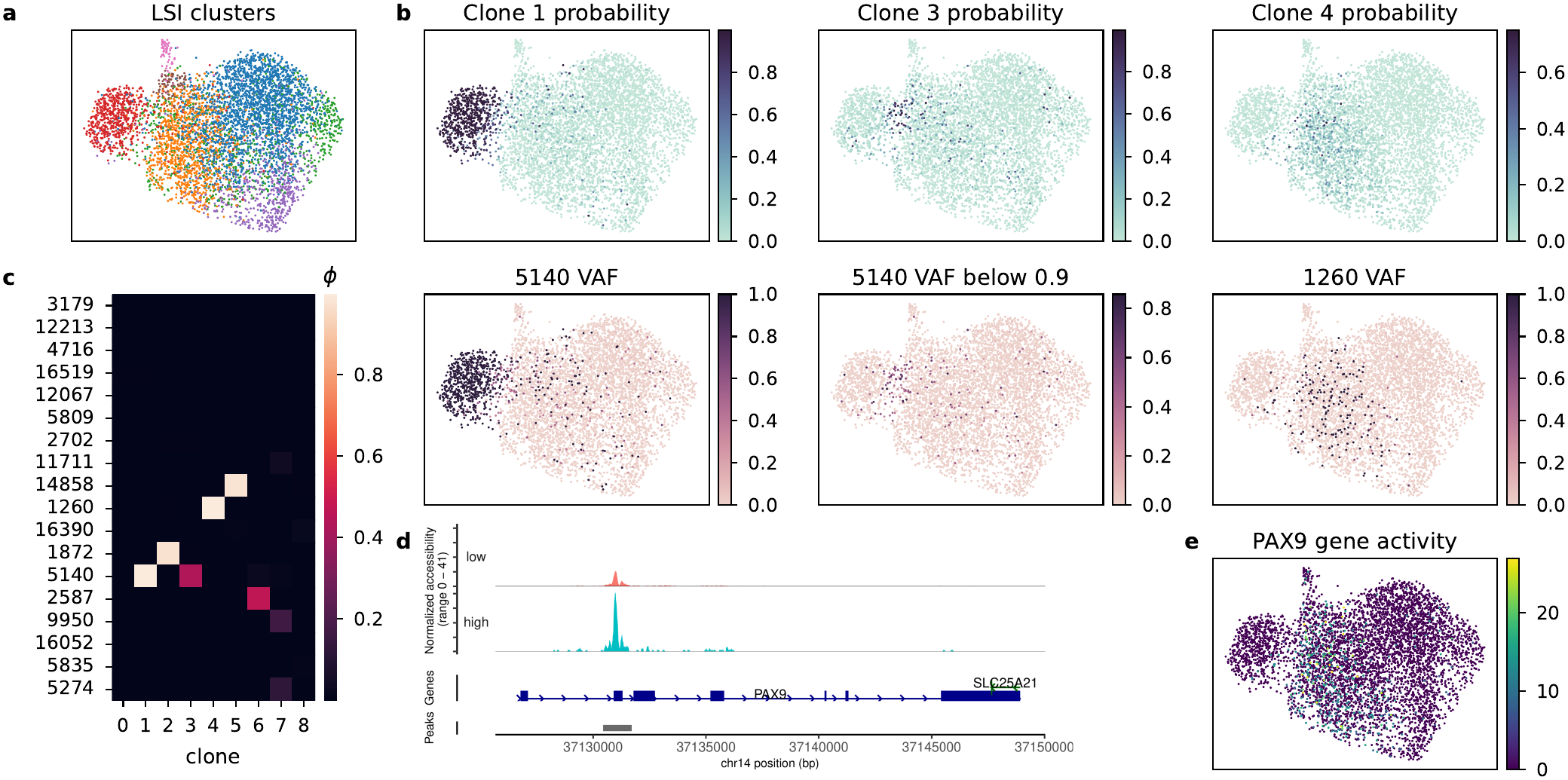
CloneVI applied to CLL mtscATAC-seq data, **a**, Clusters from latent semantic indexing (LSI) of chromatin accessibility on the UMAP of the cell-state space, **b**, Estimated probabilities for representative clones and the observed VAF of their associated mtDNA mutations; in the middle panel, VAFs above 0.9 are set to 0 to show the cells with intermediate VAF of 5140. **c**, Clone-specific allele frequencies *ϕ*. **d**, Chromatin accessibility around PAX9 in cells with high and low Clone 4 probability, **e**, PAX9 gene activity on the UMAP of the cell-state space.

Clone 1 and Clone 3 were both associated with the mtDNA mutation at position 5140, but the two populations showed different VAFs of this mutation and occupied different regions of the cell-state space (Fig. 4b). The clone-specific allele frequencies *ϕ* reflected these differences in heteroplasmy (Fig. 4c). CloneVI therefore separated two populations carrying the same mutation by their heteroplasmy levels.

We next characterized the regulatory features associated with the inferred clones. Cells with high probability of Clone 4, which carried the variant at position 1260 (Fig. 4b,c), showed increased accessibility around the PAX9 locus compared with cells with low Clone 4 probability, which also appeared as high PAX9 gene activity in the region of the cell-state space associated with Clone 4 (Fig. 4d,e). Among the genes with the highest gene activity weighted by the Clone 4 probability were also several replication-dependent histone genes (Supplementary Fig. S4d). PAX9 expression has previously been reported as a potential prognostic marker in CLL(22). Thus, CloneVI linked an mtDNA-associatcd clone with a chromatin feature relevant to the disease.

Together, these results indicate that CloneVI applies to chromatin accessibility as it does to gene expression: it separates populations that carry the same mutation at different heteroplasmy levels and relates the inferred clones to chromatin features of their cell states.

### CloneVI captures and characterizes cell-state-enriched mutant-cell populations in a large multi-donor cohort

We then applied CloneVI to an ovarian tumor mtscATAC-seq cohort(23) of 108,260 cells from five donors and several cell types (Fig. 5a,b), in which we called 64 high-confidence mtDNA variants (Methods; Supplementary Table 1). The donor was given to the model as a covariate (Methods). CloneVI inferred 17 clones (Fig. 5d; Supplementary Figs. S5 and S6). The variant 310T>C, which is carried by almost every covered cell, was largely absorbed into the shared component of the allele frequency as a background variant (Supplementary Fig. S7). Most clones were confined to a single donor: 10 of the 17 had more than 90% of their clonal probability in one donor (Supplementary Fig. S8).

**Fig. 5.**
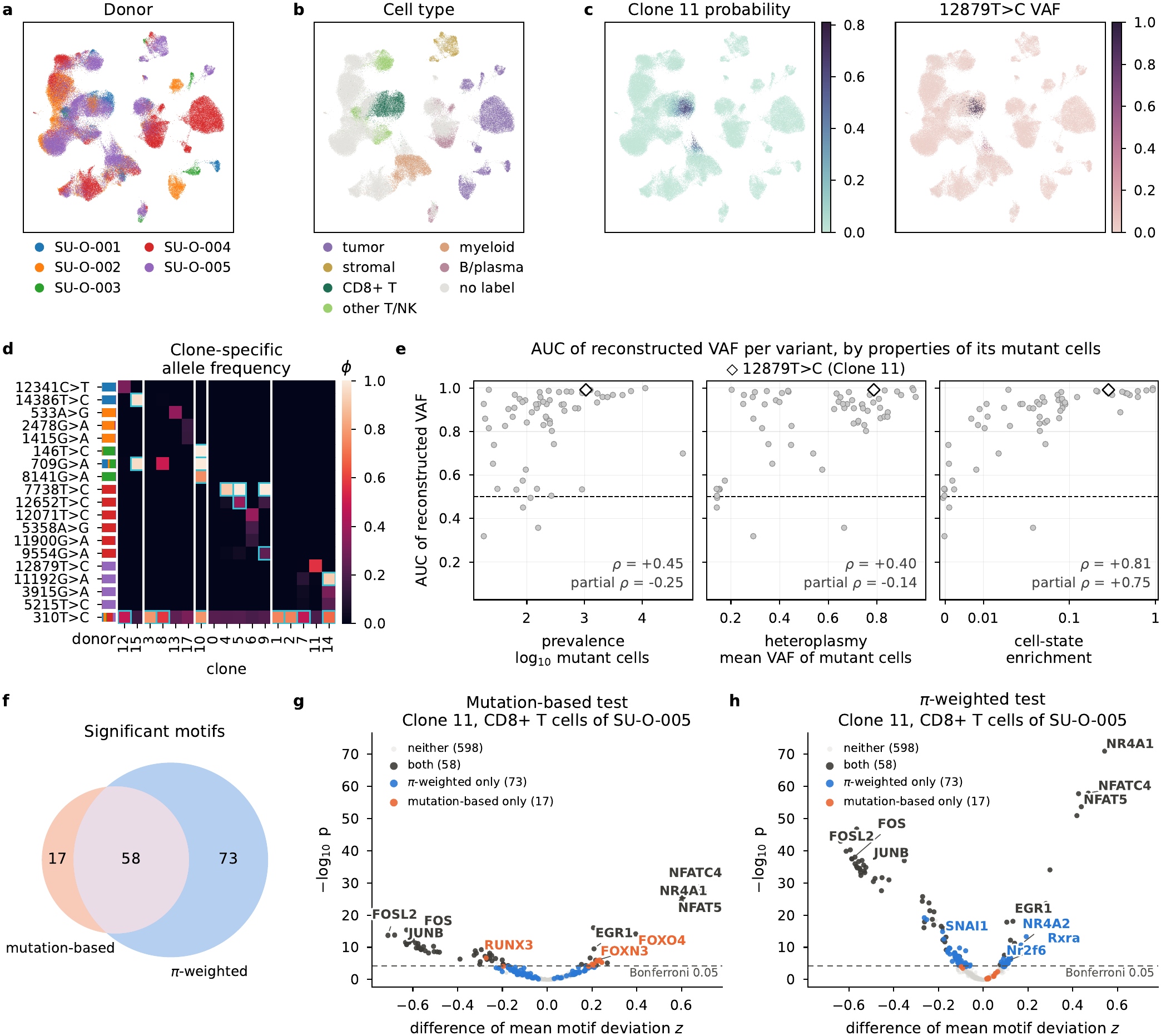
CloneVI applied to an ovarian tumor mtscATAC-seq cohort. **a–c**, UMAP of the cell-state space (108,260 cells) colored by donor (**a**), cell type (**b**; annotations of the source study, where available), and the clonal probability of Clone 11 and the observed VAF of its variant 12879T>C (**c**; gray, mtDNA depth below 10). **d**, Allele frequency ***ϕ*** of the inferred clones for 19 of the 20 variants with *ϕ* ≥ 0.1 in some clone, ordered by donor; 515A>G is not drawn (Methods). The left strip gives the donor composition of each variant’s mutant cells (colors as in **a**), and white lines separate donors. Outlined, *ϕ* at least 0.8 times the mean VAF of the variant’s mutant cells. 310T>C (bottom row) is carried by almost every covered cell. The observed VAF of each of these variants is shown in Supplementary Fig. S6. **e**, AUC with which the reconstructed VAF ***πϕ*** ranks mutant above non-mutant cells for the 64 variants, against prevalence, mean VAF of mutant cells and cell-state enrichment; *ρ*, Spearman correlation; partial *ρ*, partial Spearman correlation adjusted for the other two properties; open diamond, 12879T>C (values in Supplementary Table 1). **f–h**, Differential motif activity (chromVAR) for Clone 11 in 2,077 CD8^+^ T cells of one library of donor SU-O-005 (Supplementary Fig. S9). **f**, Motifs significant (Bonferroni-corrected *p <* 0.05) in each test, colored as in **g** and **h. g**,**h**, Volcano plots of the 746 motifs for the mutation-based test (**g**; mutant, VAF ≥ 0.10) and the ***π***-weighted test (**h**); dashed line, Bonferroni threshold.

Within a single donor, CloneVI also separated nested clones (Fig. 5d). Clones 4, 5 and 9 of donor SU-O-004 all carried 7738T>C, and Clones 5 and 9 were distinguished by additional variants: 12652T>C, carried mainly by Clone 5, and 9554G>A, a variant observed at a mean VAF of 0.25 and carried only by Clone 9 (Supplementary Fig. S8c–e).

As in the MAESTER data, the AUC was positively correlated with cell-state enrichment (*ρ* = +0.81; partial *ρ* = +0.75; Fig. 5e). Prevalence and mean VAF were also correlated with the AUC (*ρ* = +0.45 and +0.40), but they were themselves correlated with enrichment (*ρ* = +0.69 and +0.57), and after adjustment their partial correlations were small and negative (partial *ρ* = −0.25 and −0.14). For example, 3109T>C was mutant in 902 cells across all five donors, but mostly at low VAF (above 0.50 in only one cell); it showed no cell-state enrichment and was not captured (AUC 0.53; Supplementary Fig. S9d). Because the donor is also an input of the decoder for ***π***, a variant confined to one donor could appear enriched and be captured through the donor alone. When enrichment and AUC were recomputed within each donor, the AUC remained positively correlated with enrichment in every donor (*ρ* = +0.10 to +0.68; Supplementary Fig. S10).

Next, to see how a clone differs from a mutant-cell population, we used each of them to define the groups of a differential test of transcription factor motif activity. The clonal probability is estimated from the cell state and varies continuously, whereas each cell is called mutant or non-mutant from its observed VAF. We chose Clone 11, which carries 12879T>C and is found in CD8^+^ T cells of donor SU-O-005 (Fig. 5b,c; Supplementary Fig. S9). Motif activity was computed with chromVAR(24), which scores the accessibility of the binding sites of each transcription factor in each cell. On the same 2,077 CD8^+^ T cells of one library of this donor (Methods), we performed a mutation-based test, which compares mutant with non-mutant cells, and a ***π***-weighted test, which weights each cell by its clonal probability. Because the clonal probability is itself estimated from chromatin accessibility, the ***π***-weighted test describes the state in which the cells of the clone lie; it does not test whether the variant itself changes motif activity. Of the 746 motifs, 75 were significant in the mutation-based test and 131 in the ***π***-weighted test, with 58 in both (Fig. 5f). No motif significant in either test changed in opposite directions, and the effect sizes of the two tests correlated at 0.93 over all motifs (Fig. 5g,h).

The 58 motifs found by both tests showed increased activity of NFAT-family, NR4A1 and EGR motifs and decreased activity of AP-1 motifs. This combination has been described in CD8^+^ T cells under chronic antigen stimulation, in which NFAT acting without its partner AP-1 induces NR4A and EGR2 and establishes an exhaustion program(25–27).

The 73 motifs found only by the ***π***-weighted test had smaller effects than the shared motifs, with the same sign and similar size in the mutation-based test; they included NR4A2, another member of the NR4A family. The 17 motifs found only by the mutation-based test involved RUNX3 and FOX factors rather than the exhaustion circuit.

Together, these results indicate that, in this cohort, CloneVI captures mutant-cell populations according to their cell-state enrichment rather than their prevalence or mean VAF, and that weighting by the clonal probability characterizes the cell state in which the mutant cells concentrate.

## Discussion

CloneVI integrates mtDNA mutation information with cell states measured by single-cell RNA-seq or ATAC-seq. Whereas existing approaches first fix clone labels using genotype alone, CloneVI estimates clonal probabilities from the cell state while estimating the mutations of each clone. The analysis therefore focuses on genetic clones that are linked to cell states.

CloneVI is designed to capture mutant-cell populations that are associated with cell states. A genetic clone whose mutant cells are not enriched in particular states of gene expression or chromatin accessibility is therefore not captured as a clone, even though its mutant cells are present in the data. Such genetic clones are the target of genotype-first methods. The two approaches are complementary: genotype-first methods describe the genetic clonal structure itself, whereas CloneVI identifies which parts of that structure are associated with cell states. For the same reason, the clonal probability describes how a clone is distributed over cell states rather than the genotype of each cell, and a differential test weighted by it characterizes the cell states of the clone.

In this study, we used mtDNA mutations as genetic information. Because clonal probabilities are estimated from cell state, they are defined for every cell, including cells in which a variant is not covered. This property may allow CloneVI to be extended to nuclear somatic mutations detected in scRNA-seq or scATAC-seq data, whose detection is limited by dropout and low coverage.

## Conclusion

CloneVI provides a probabilistic framework for integrating cell states with mtDNA mutations. By estimating clonal probabilities from gene expression or chromatin accessibility while reconstructing mutation counts, the method identifies genetic clones associated with cell-state diversity and characterizes their molecular properties. CloneVI is applicable to both scRNA-seq and scATAC-seq data and provides a basis for studying genetic and non-genetic heterogeneity together at single-cell resolution.

## Methods

### Model

For simplicity, we describe the model at the level of a single cell *c*. Throughout, *c* indexes cells, *k* clones and *m* mtDNA variants. The graphical model is shown in Supplementary Fig. S1.

#### Cell state inference

CloneVI is based on a variational autoencoder (VAE)(21), which compresses the input data and reconstructs it to infer a latent cell state. The input is a count vector of *J* features, such as gene expression or chromatin accessibility, denoted as **x**_*c*_ *∈*ℕ^*J*^. An encoder with parameters **Φ** maps **x**_*c*_ to the parameters of a normal distribution, ***µ***_*c*_ *∈* ℝ^*L*^ and 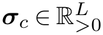, from which the cell state **z**_*c*_ *∈* ℝ^*L*^ is sampled:

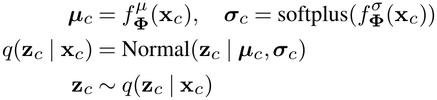

The latent dimension was set to *L* = max(10, *J* ^1*/*4^) and the hidden dimension to max(50, ⌊ *J* ^1*/*2^ ⌋), where ⌊· ⌋ denotes rounding down to an integer.

The input counts are reconstructed from the cell state by a decoder with parameters **Θ**_*x*_, which outputs the mean ***λ***_*c*_ of a negative binomial distribution with a trainable feature-wise dispersion 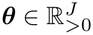. The mean is the count expected from the totals of the cell and the feature, multiplied by a positive factor that the decoder outputs for each feature:

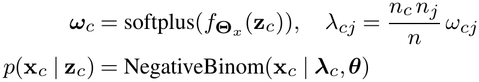

where *n*_*c*_ and *n*_*j*_ are the total counts of cell *c* and feature *j*, and *n* is the total count of the matrix.

#### Estimation of clonal probability based on cell state

From the cell state, the clonal probability ***π***_*c*_(**z**_*c*_) *∈* [0, 1]^*K*^ is estimated by a decoder with parameters **Θ**_*b*_, where *K* is the number of clones:

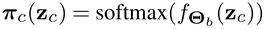

We write *π*_*ck*_ for the *k*-th element of ***π***_*c*_(**z**_*c*_). When the data contain several samples, a one-hot indicator of the sample of origin is concatenated to the input of the encoder and to the inputs of both decoders, so that the decoder for **x**_*c*_ can explain variation specific to a sample without **z**, and the decoder for ***π*** can use the sample in predicting the mutations.

To estimate ***π***_*c*_(**z**_*c*_), we maximize the likelihood of the counts of *M* mtDNA mutations **b**_*c*_ ℕ^*M*^ observed in each cell, modeled with a binomial distribution. The clone *ρ*_*c*_ of cell *c* follows a categorical distribution with the clonal probability, and if *ρ*_*c*_ = *k*, the mutation counts are observed with the clone-specific allele frequency ***ϕ***_*k*_ *∈* [0, 1]^*M*^, the expected VAF of each variant in clone *k*, and the sequencing depth **d**_*c*_ *∈* ℕ^*M*^:

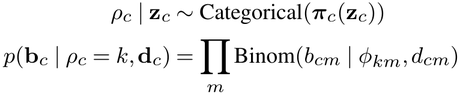

Each cell is therefore not assigned deterministically to a single clone, and marginalizing *ρ*_*c*_ gives a mixture weighted by the clonal probability:

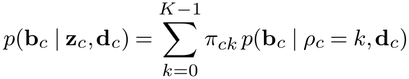

Optionally, the allele frequency of each clone is decomposed into an allele frequency shared by all clones, ***ϕ***^(common)^ *∈* [0, 1]^*M*^, and a clone-specific component 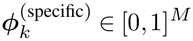:

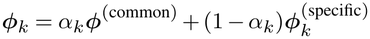

where *α*_*k*_ *∈* [0, 1] keeps ***ϕ***_*k*_ within [0, 1]. 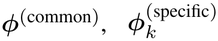 and *α*_*k*_ are trainable parameters. Whether this decomposition and the other options below were used in each analysis is listed in Supplementary Table 2.

#### Loss function

The model is trained by minimizing

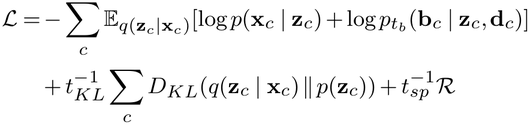

where *p*(**z**_*c*_) is a standard normal prior, ℛ is an optional regularization term, and *t*_*KL*_, *t*_*b*_ and *t*_*sp*_ are temperatures. When all temperatures equal 1 and ℛ is omitted, ℒ is the negative evidence lower bound (ELBO) of the log marginal likelihood of the observed data.

A larger number of clones allows the allele frequencies to be fitted more flexibly and can lead to unnecessary clones. The regularization term, based on group lasso(28), therefore increases the loss when many clones are used:

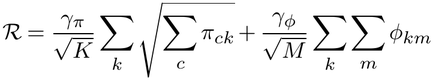

The temperatures start large and decrease to 1, so that the weights of their terms increase gradually. The temperature *t*_*KL*_ prevents collapse of the latent variables, and *t*_*b*_ and *t*_*sp*_ prevent the clonal probabilities from collapsing onto a single clone early in training. The temperature *t*_*b*_ is applied to each term of the mixture:

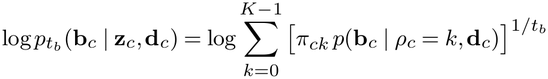

While *t*_*b*_ is large, this lowers the weight of the mutation likelihood and evens out the contributions of the clones; at *t*_*b*_ = 1 it equals log *p*(**b**_*c*_ | **z**_*c*_, **d**_*c*_).

The loss is minimized in stages. First, the encoder and decoder are trained on the reconstruction of **x**_*c*_ and the KL divergence. Then, with the encoder, the decoder for **x**_*c*_ and the dispersion ***θ*** fixed, the decoder for ***π*** and the allele frequencies ***ϕ*** are estimated from the mutation likelihood. When regularization is used, ℛ is added in a final stage. Cells were split into training (85%), validation (5%) and test (10%) sets. Each temperature decreases to 1 during the stage in which its term is first trained (*t*_*KL*_ in the first stage, *t*_*b*_ in the second and *t*_*sp*_ in the third), and a stage can end only after its temperature has reached 1 and, in the first stage, after a warm-up in which the learning rate is increased. A stage then ends when its loss on the validation set, computed without temperatures, stops improving, and the parameters with the lowest such loss during the stage are restored (Supplementary Algorithm 1). Because the encoder is fixed after the first stage, the cell state is inferred from **x**_*c*_ alone and does not use mtDNA mutation information. The settings of each analysis are listed in Supplementary Table 2.

#### Pruning of clones

The number of clones *K* is a user-specified upper bound; we used *K* = 20 for the real datasets and *K* = 10 for the simulation (Supplementary Table 2). After training, clones carrying little probability were removed to make the clones easier to interpret: clone *k* was retained if 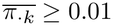 or max_*c*_ *π*_*ck*_ *≥* 0.2, and other clones were removed together with their allele frequencies. All clones were kept for the comparison of prediction accuracy (see below).

### Preparation of datasets and downstream analysis

#### Generation of simulation data

Based on the Scanpy PBMC3k dataset(29), we simulated a scenario in which each cell belongs to one of two clones and carries the corresponding mutation profile. For each of the eight clusters in the dataset, a clonal probability ***π*** *∈* [0, 1]^2^ shared by all cells in the cluster was set: fixed for three clusters and drawn at random for the other five. Each cell was assigned to a clone *ρ*_*c*_ *∈ {*0, 1*}* by a Bernoulli draw with this probability. The sequencing depth **d**_*c*_ *∈* ℕ^*M*^ at each of *M* loci was drawn from Poisson(80). A binary mutation profile **r**_*k*_ *∈ {*0, 1*}* ^*M*^ was defined for each clone, and the mutation counts were calculated as **b**_*c*_ = **r**_*ρc*_ ⊙ **d**_*c*_, where ⊙ denotes the Hadamard product. The simulation settings and the clonal probabilities of the eight clusters are listed in Supplementary Table 3, and the settings used to fit CloneVI in Supplementary Table 2; the generated data are provided with the code.

#### Processing of MAESTER and CLL mutation data

For the MAESTER(12) and CLL mtscATAC-seq(11) datasets, the mutation calls of maegatk(12) and mgatk(11) provided with the original studies were used. For each locus, the counts of bases other than the reference allele were summed to define the mutation count, so the variants of these two datasets are named by their position. The variants analyzed in the original studies were used: 26 for the MAESTER data and 18 for the CLL data. The variants of the ovarian cohort were called per allele (see below).

#### Preprocessing of MAESTER gene expression data

MAESTER data of bone marrow cells from patient BPDCN712 were obtained from GSE182685(12). The rds files were converted to h5ad files and processed with Scanpy(29). Cells with at least 200 detected genes and genes detected in at least three cells were retained. Counts were normalized, and the raw counts of the top 2,000 highly variable genes were used for model training. For downstream analysis, genes detected in at least 20 cells were retained, counts were log-normalized, and the top 4,000 highly variable genes were used, each divided by its mean over cells.

#### Downstream analysis of MAESTER data

Clone-associated expression was summarized for each gene and clone as the ***π***-weighted mean expression: the mean over cells of the log-normalized expression of the gene, divided by its mean over cells, with each cell weighted by its probability *π*_*ck*_ of the clone. For each displayed clone, the TCR variable-region gene with the highest ***π***-weighted mean expression was shown on the UMAP of the cell-state space.

The reconstructed VAF of variant *m* in cell *c*, written ***πϕ*** in the Results and figures, is the VAF expected under the fitted mixture:

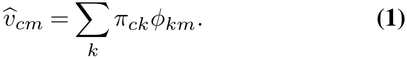

In the test cells, which were not used for training, the reconstructed VAF was compared with a baseline that assigned each cell the mean observed VAF of its annotated cell type in the cells outside the test set. Agreement with the observed VAF was evaluated for each variant by Pearson correlation. The baseline was computed outside the test set because cell-type means are taken directly from observed VAFs and would otherwise include the evaluated cells. Because this comparison evaluates prediction accuracy, the reconstruction used all *K* clones of the fitted model, including those removed by pruning.

#### Cell-state enrichment and capture of mtDNA variants

For each variant *m*, a cell was defined as covered if its mtDNA depth was at least 10, and as mutant if, in addition, its VAF was at least 0.10 and at least three reads carried the alternate allele; let *C*_*m*_ be the covered cells and *A*_*m*_ *⊆ C*_*m*_ the mutant cells. Variants with at least 15 mutant and 15 non-mutant cells were analyzed.

Cell-state enrichment was computed in the cell-state space without using the clonal probabilities. On a 30-nearest-neighbor graph of all cells (Euclidean distance on standardized **z**), with *N*_*c*_ the neighbors of cell *c*, the local purity of the mutant cells, the mean fraction of the neighbors of a mutant cell that are also mutant, is

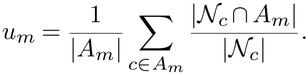

Because purity also increases with the number of mutant cells, we estimated the purity expected by chance by permuting the mutant labels 200 times among the covered cells *C*_*m*_; with 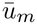 the mean purity over the permutations, enrichment was defined as

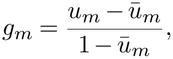

which is 0 for randomly placed mutant cells and 1 when all neighbors of mutant cells are mutant.

Capture of a variant by the model was measured as 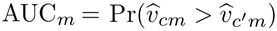 for *c A*_*m*_ and *c*^*′*^ *∈C*_*m*_ *\ A*_*m*_, the probability that a mutant cell receives a higher reconstructed VAF Eq. (1) than a non-mutant cell; the reconstruction used only the clones retained by pruning. Across variants, enrichment, prevalence (log_10_ |*A*_*m*_|) and mean VAF of mutant cells were each related to AUC_*m*_ by Spearman correlation and by partial Spearman correlation adjusted for the other two properties. Because variants share cells, these correlations are reported without *p*-values. As a sensitivity analysis, removing the requirement of three alternate reads left the partial correlation of enrichment positive (+0.80 in the ovarian cohort and +0.57 in the MAESTER data) but the partial correlations of prevalence and mean VAF were then positive in both datasets.

#### Preprocessing of CLL mtscATAC-seq chromatin accessibility data

CLL mtscATAC-seq data were obtained from the original study(11). Output data from cellranger-atac were loaded and processed with Signac(30). After constructing the cell-by-peak fragment count matrix, peaks observed in at least 10 cells and cells with at least 200 detected peaks were retained, and their raw fragment counts were used as the model input. The data were converted from an rds file to an h5ad file and processed with Scanpy(29).

#### Downstream analysis of CLL mtscATAC-seq data

Gene activity was calculated with the GeneActivity function of Signac(30), normalized to 10,000 counts per cell and log-transformed; the 2,000 most variable genes were kept and each gene was divided by its mean over cells. For each clone, genes were ranked by their ***π***-weighted mean gene activity, computed in the same way as for the MAESTER data; PAX9 ranked first for Clone 4. For each clone, cells with high clonal probability were defined as the cells with the largest *π*_*ck*_ that together accounted for 50% of the total probability of the clone, and the remaining cells were defined as cells with low clonal probability; accessibility in the two groups was shown with CoveragePlot. The adjusted *p*-value printed on the coverage plot of Supplementary Fig. S4d is from the Wilcoxon rank-sum test of gene activity between the two groups (Find-Markers in Seurat(31), Bonferroni correction over all genes) and is descriptive only, because the gene was selected by the ranking above and the two groups are defined from the same accessibility data.

#### Preprocessing of ovarian mtscATAC-seq data

The ovarian tumor mtscATAC-seq cohort was obtained from GSE302113(23). Because the deposited tables do not contain the per-cell mitochondrial coverage that the binomial likelihood requires, the raw libraries were reprocessed with cellranger-atac 2.1.0 against GRCh38, mitochondrial allele counts were obtained with mgatk 0.7.0(11), and every library was recounted against a merged peak set with Signac(30). Cells were retained with 1,000–50,000 peak counts, more than 40% of reads in peaks, a blacklist ratio below 0.05, a nucleosome signal below 4 and a TSS enrichment above 3, leaving 108,260 cells from five donors; the raw counts of the top 25% of peaks (85,859) were used as the cell-state input. High-confidence variants were called per donor following the Signac mitochondrial vignette, requiring confident detection in at least max(5, 0.005*N*_donor_) cells, giving 64 variants in the union. One of them, 515A>G, was called only in donors SU-O-001 and SU-O-002, in which its position is covered in far fewer cells than in the other donors and than the neighboring positions, so it is unlikely to mark a population of cells; it is kept in all analyses but not drawn in Fig. 5d. Cell-type annotations were taken from the source study and matched by barcode; they were available for 45,678 cells in 7 of the 11 libraries and were grouped into six classes for display. The donor (five levels) was given to the model as the indicator of the sample of origin.

#### Within-donor analysis of the ovarian cohort

For the within-donor analysis (Supplementary Fig. S10), enrichment and AUC were recomputed in each donor using only its covered cells: the neighbor graph was built on these cells with **z** standardized within the donor, the non-mutant cells were those of the same donor, and the mutant labels were permuted 500 times.

#### Motif activity and clone-probability-weighted differential testing

Motif activity was computed once for the whole cohort with chromVAR(24) as implemented in Signac, using the 746 vertebrate CORE motifs of JASPAR2020(32), and is reported as the deviation *z* score of each motif in each cell. For this test a cell was called mutant at an mtDNA depth ofat least 10 and a VAF of at least 0.10. The requirement of three alternate reads used in the analysis of cell-state enrichment was not applied here; it would have moved 13 of the 634 mutant cells, whose median VAF is 1.00, to the group of 1,443 non-mutant cells. The mutation-based test was the Wilcoxon rank-sum test of Seurat(31) applied to this assay, with the difference of group means as the effect size, no fold-change filter and Bonferroni correction over motifs. In the ***π***-weighted test, each cell *c* is counted in both groups, as a fraction *w*_*c*_ = *π*_*ck*_ of a cell in the clone group and 1 − *w*_*c*_ in the reference group, and the same Wilcoxon test is computed with these weights; with weights of only 0 and 1 it gives the same *p*-values as the Seurat function. When the clonal probabilities were permuted across cells, 5% of motifs had *p <* 0.05, as expected when there is no difference.

Cells for both tests were the annotated CD8^+^ T cells of the library containing most of the clonal probability of the clone, covered at depth at least 10 at the variant position, giving 634 mutant and 1,443 non-mutant cells for Clone 11. Because cells are not independent observations, the thresholds are common reference points rather than calibrated error rates.

### Software

The software and versions used in this study are listed in Supplementary Table 4; the training settings of each fit are provided with the code.

### Use of generative AI tools

Claude (Anthropic) was used to write preprocessing, analysis and plotting code, mainly for the ovarian tumor cohort, and to reorganize the analysis and plotting code for the other datasets for public release. The CloneVI source code was written by the authors and later modified and reorganized using Claude. Claude was also used to turn the authors’ outlines and drafts into manuscript text, adding the descriptions of the analysis results; the authors then revised and edited this text. Codex (OpenAI) was used to review analysis code, and Gemini (Google) for partial language editing of the initial manuscript. The authors reviewed all code, results and text and take full responsibility for the content of this article.

## Author contributions statement

Y. Kojima conceived the idea for this study. T. Hibi designed and performed the experiments under the supervision of Y. Kojima and T. Shimamura. S. Hayashi and C. Mizukoshi advised on the implementation, and H. Hirose and C. Mizukoshi advised on the manuscript. All authors have read and approved the manuscript.

## Data availability

The datasets used in this study are publicly available. The Scanpy PBMC3k dataset is available from 10x Genomics, the MAESTER dataset from GSE182685, the CLL mtscATAC-seq dataset from https://osf.io/bupge/, and the ovarian tumor mtscATAC-seq dataset from GSE302113. The source code of CloneVI is available at https://github.com/TaichiHIBI/cloneVI. The simulated data, the processed inputs and the fitted models analyzed in this study are deposited at Zenodo (https://doi.org/10.5281/zenodo.23687068), and the code that reproduces the figures and numbers of this article from them is included in the repository.

## Acknowledgments

Taichi Hibi would like to thank the THERS Nagoya University Program for Fostering Next Generation AI Professionals. The use of generative AI tools is described in Methods.

## Alt text for figures

Figure 1. Alt text: Schematic in two parts. The top part contrasts an existing approach, which assigns cells to clones from mutations alone, with CloneVI, which gives each cell probabilities of belonging to each clone within a cell-state space. The bottom part shows the model: non-genetic data x are encoded into a latent cell state z and decoded back, and mtDNA mutation counts b are reconstructed from z through the clonal probabilities *π* and the clone-specific allele frequencies *ϕ*, both drawn as heatmaps.

Figure 2. Alt text: Panel a shows a UMAP of simulated PBMC clusters labeled with clone mixing ratios and a diagram of how mutation counts were simulated. Panel b shows three UMAPs, the true clone labels and the estimated probabilities of clones 0 and 1, whose high-probability regions match the true clone areas. Panel c is a matrix of the clusters with one clone or both against summed estimated probabilities with most mass on the diagonal, and panels d and e are bar charts of nearly identical true and estimated clone proportions, about 64% and 36%.

Figure 3. Alt text: Panel a is a UMAP of bone marrow cells colored by cell type, with T cells and cytotoxic T lymphocytes labeled. Panel b is a heatmap of allele frequencies for 26 variants across the inferred clones, where a few bright cells mark clone-specific variants. Panel c shows UMAPs of the probabilities of three clones above UMAPs of the observed VAF of their variants, which highlight the same regions, and panel d shows expression of three T-cell receptor genes: TRBV7-9 and TRDV2 are concentrated in the regions of Clones 3 and 5, and TRDV1 is scattered over the CTL region of Clone 1. Panel e is a scatter plot comparing per-variant VAF correlations of CloneVI with a cell-type baseline, and panel f plots AUC against prevalence, heteroplasmy and cell-state enrichment in three plots side by side, with AUC rising most clearly with enrichment.

Figure 4. Alt text: Panel a is a UMAP of CLL cells colored by cluster. Panel b shows UMAPs of the probabilities of three clones above UMAPs of the observed VAF of variants 5140 and 1260, which light up matching regions. Panel c is a heatmap of allele frequencies for 18 variants across clones, panel d is a genome track in which accessibility around PAX9 is higher in cells with high Clone 4 probability, and panel e is a UMAP of PAX9 gene activity.

Figure 5. Alt text: Panels a to c are UMAPs of 108,260 ovarian tumor cells colored by donor, by six cell-type classes with CD8+ T cells in dark green, and, in panel c, by the probability of Clone 11 and by the VAF of 12879T>C, which are high in the same CD8+ T-cell region. Panel d is a heatmap of the fitted allele frequency of 19 variants against 17 clones, grouped by donor, in which almost every clone carries one variant and the exceptions are 709G>A, carried by one clone in each of three donors, and 7738T>C, carried by three clones of one donor that are told apart by 12652T>C and 9554G>A. Panel e has three scatter plots of AUC for 64 variants against prevalence, heteroplasmy and cell-state enrichment; AUC increases most steeply with enrichment, and 12879T>C, the variant of Clone 11, sits near the top of all three. Panel f is a Venn diagram of significant motifs (17 mutation-based only, 58 shared, 73 weighted only), and panels g and h are volcano plots of motif differences from the mutation-based test and the probability-weighted test, in which NFAT, NR4A1 and EGR motifs increase and AP-1 motifs decrease in both.

## Supplementary information

**Fig. S1.**
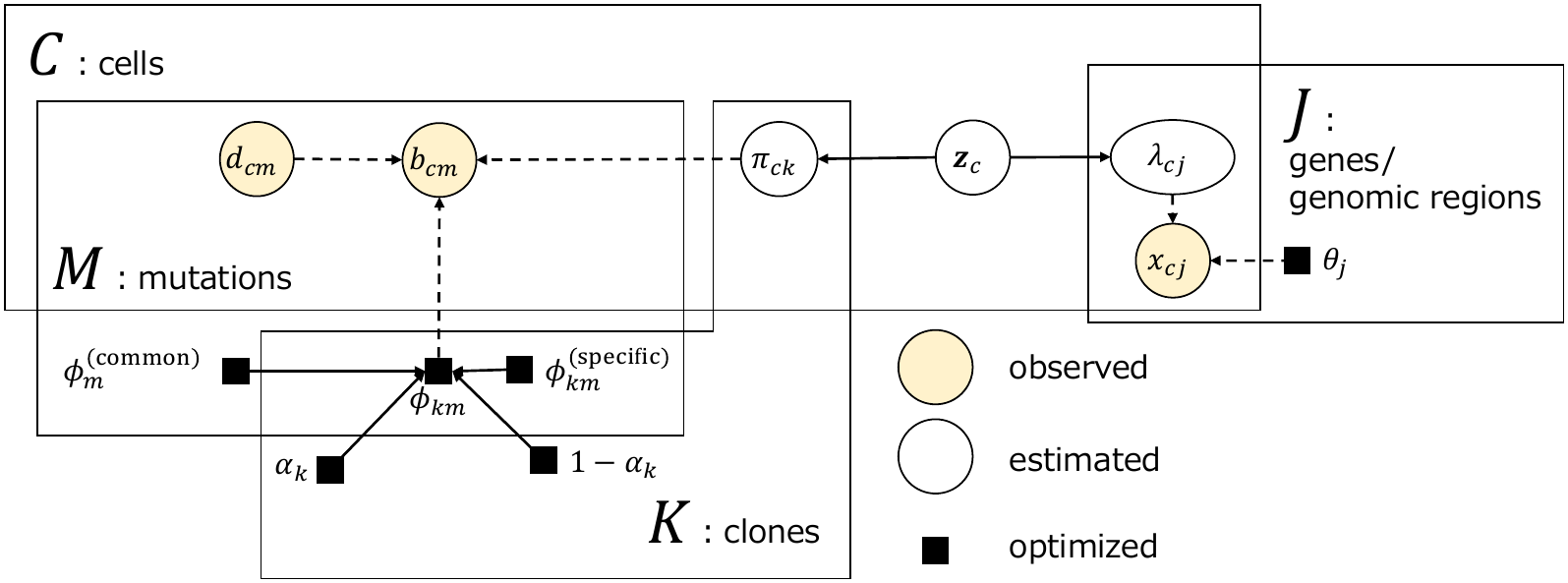
Graphical model of CloneVI. The counts **x**_*c*_ of gene expression or chromatin accessibility and the mtDNA mutation counts **b**_*c*_ of cell *c* are reconstructed from its cell state **z**_*c*_. **x**_*c*_ follows a negative binomial distribution with mean ***λ***_*c*_ and feature-wise dispersion ***θ*. b**_*c*_ follows a binomial distribution with the sequencing depth **d**_*c*_ and the allele frequency ***ϕ***_*k*_ of the clone *k* to which the cell belongs, and the likelihoods of the clones are weighted by the clonal probability *π*_*ck*_ and summed. ***ϕ***_*k*_ mixes the shared component ***ϕ***^(common)^ and the clone-specific component 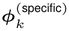 with weight *α*_*k*_ (Methods).

### Supplementary Algorithm 1

Training procedure for CloneVI. The loss is minimized in stages, and each temperature decreases to 1 during the stage in which its term is first trained. A stage can end only after its temperature has reached 1 (in Stage 1, also after the warm-up of the learning rate); it then ends when its validation loss, computed without temperatures, stops improving, and the parameters with the lowest validation loss during the stage are restored. Notation as in Methods.

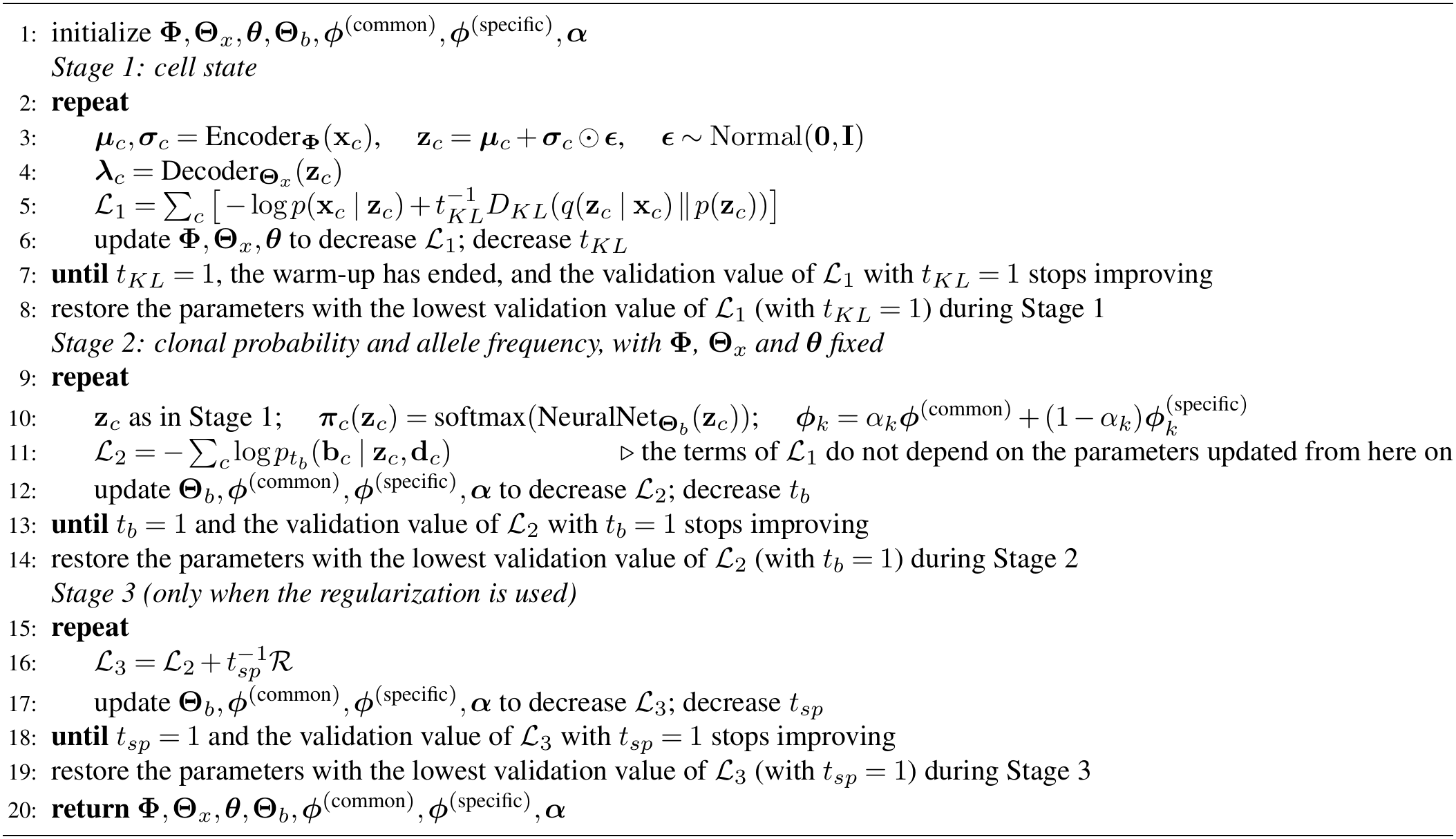

**Fig. S2.**
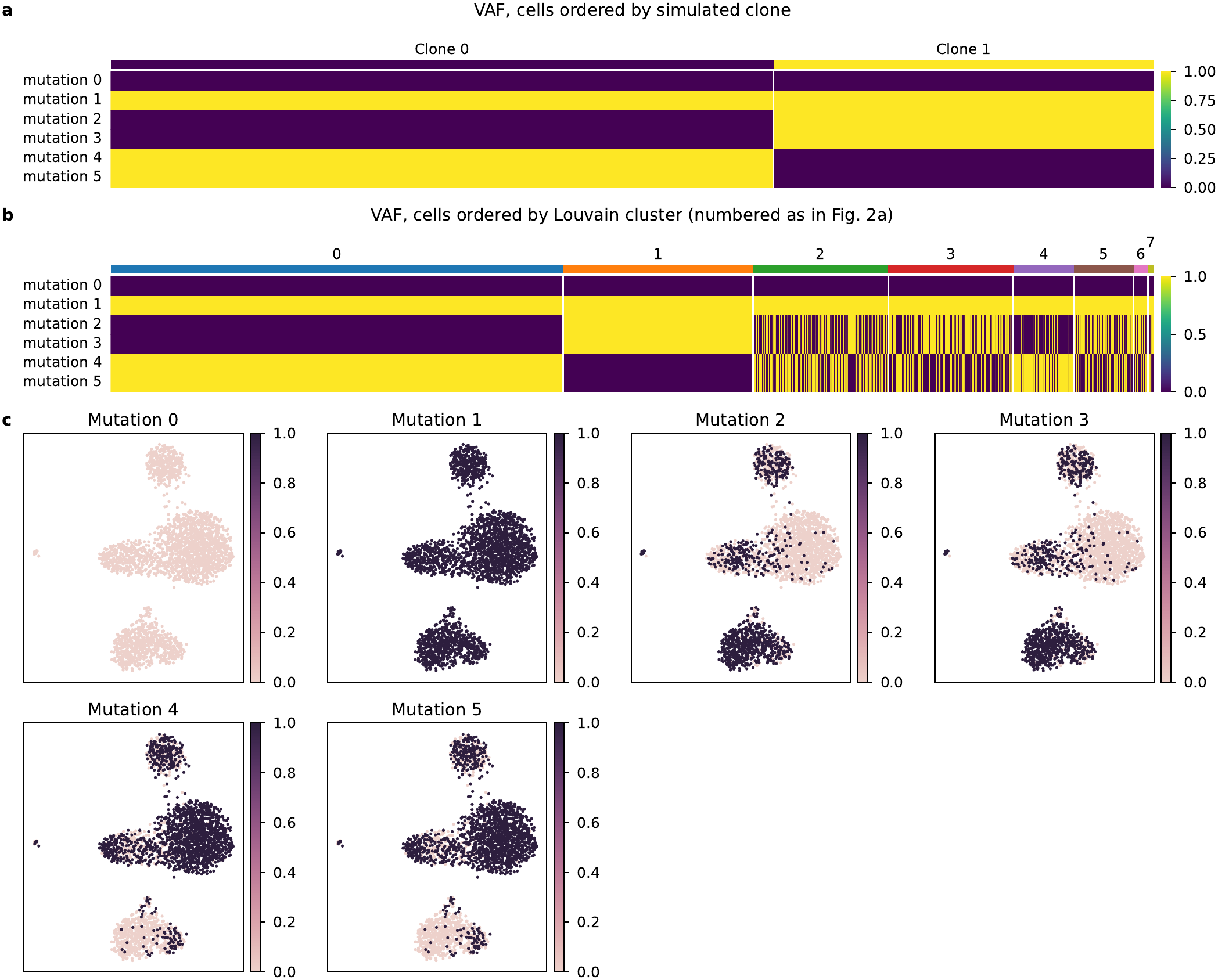
Simulated mutation data. Each cell of the Scanpy PBMC3k dataset was assigned to one of two clones with the probability specified for its Louvain cluster, and the two clones were characterized by six mutations. **a**, VAF of the six mutations (rows) in each cell, with cells ordered by clone. **b**, The same VAF with cells ordered by Louvain cluster. **c**, VAF of each mutation on the UMAP of the cell-state space.

**Fig. S3.**
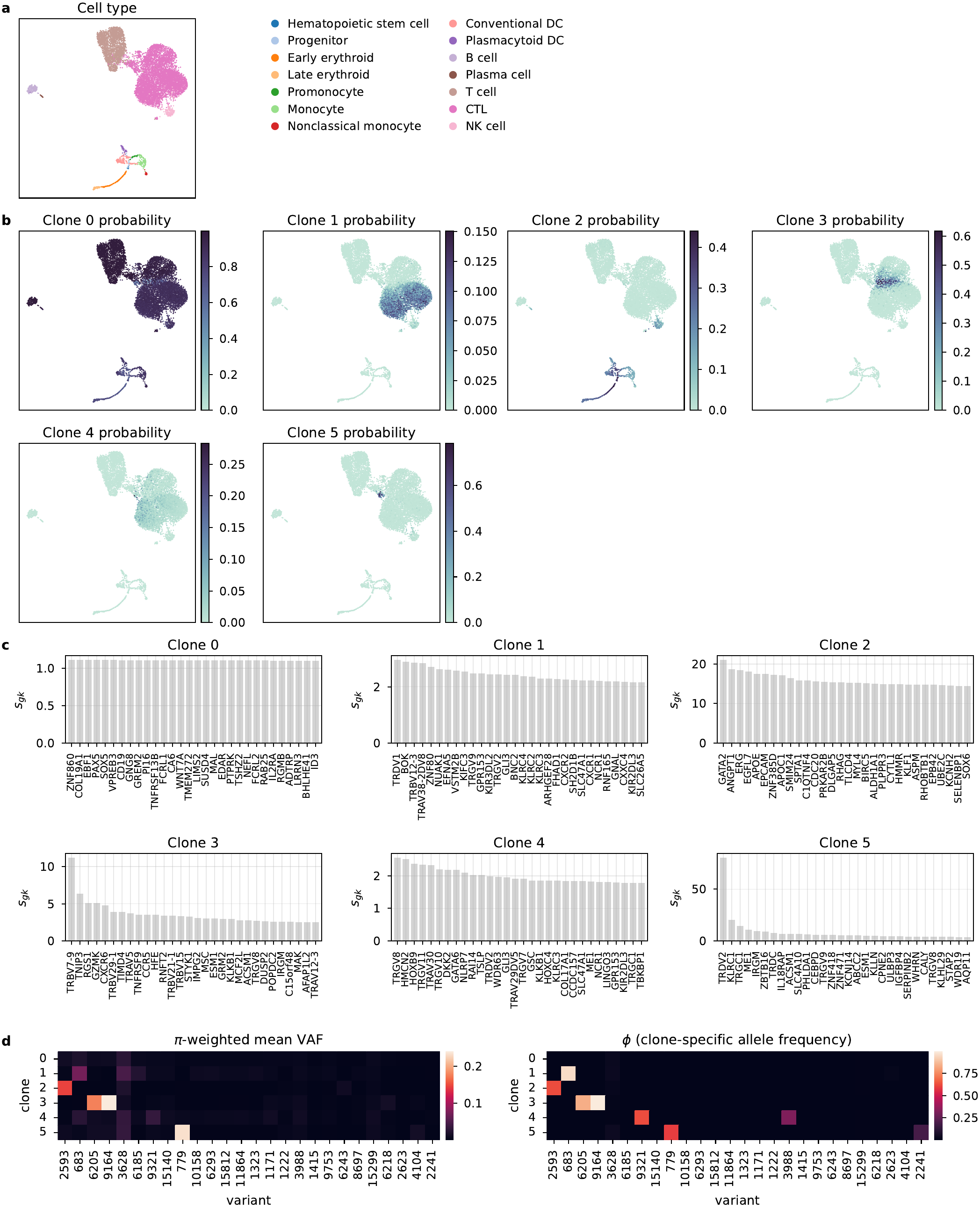
Clonal probabilities, clone-associated genes and allele frequencies in the MAESTER data. **a**, Cell types on the UMAP of the cell-state space, as in Fig. 3a. **b**, Clonal probabilities ***π*** of the six inferred clones (Clones 0–5; numbered as in Fig. 3). **c**, The 30 genes with the highest ***π***-weighted mean expression *s*_*gk*_ in each clone, the mean expression of gene *g* over cells weighted by the probability *π*_*ck*_ of clone *k* (Methods). For Clone 1, several TCR variable-region genes rank high (TRDV1, TRBV12-3 and TRAV38-2DV8), but none stands out, unlike TRBV7-9 for Clone 3 and TRDV2 for Clone 5. **d, *π***-weighted mean of the observed VAF (left) and the clone-specific allele frequency ***ϕ*** (right) for each clone and variant.

**Fig. S4.**
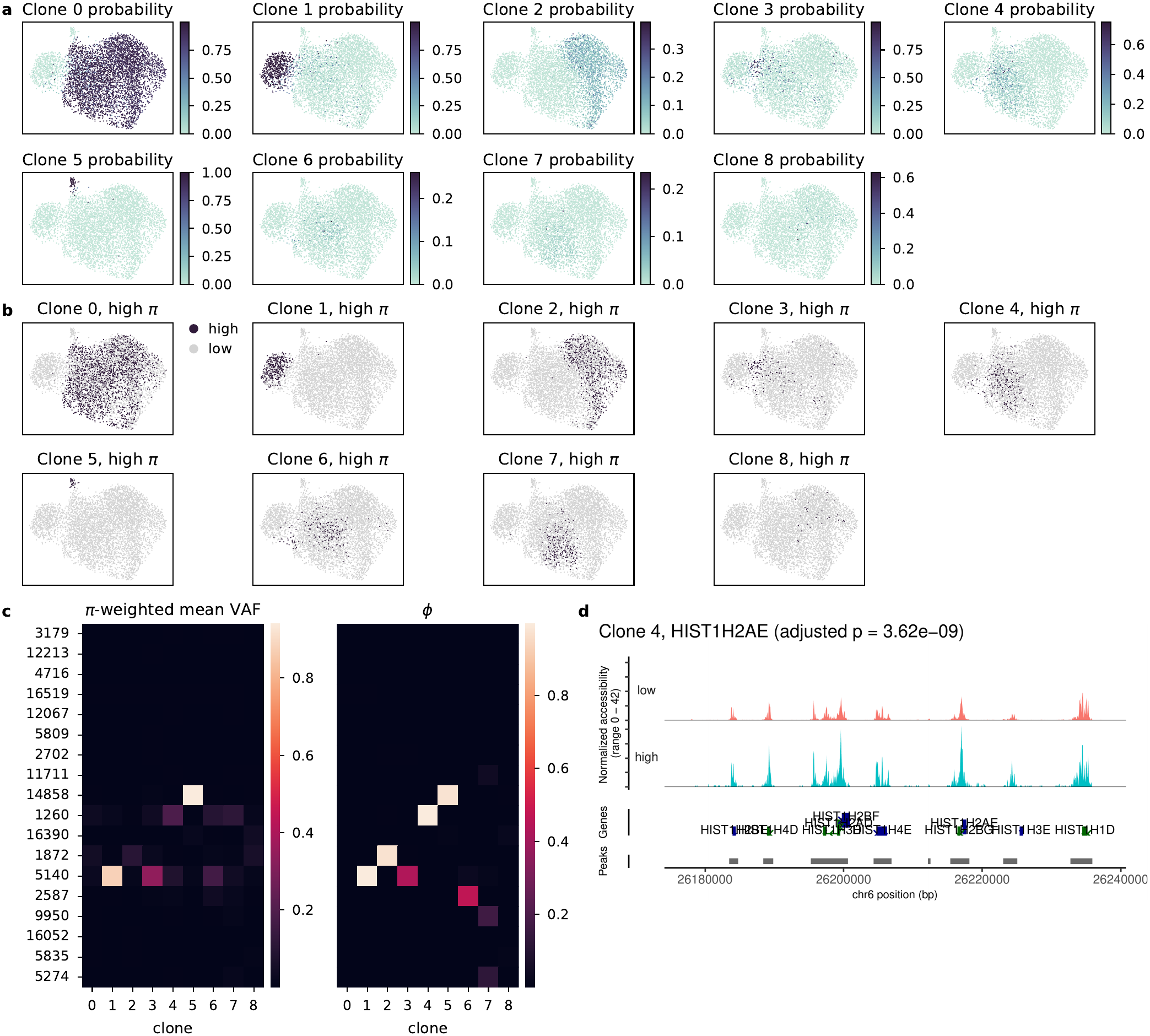
Clonal probabilities and allele frequencies in the CLL mtscATAC-seq data. **a**, Clonal probabilities ***π*** of the nine inferred clones, **b**, Cells with high probability for each clone (the cells with the largest tv that together account for 50% of the clone’s probability), used to group the cells in **d. c, *π***-weighted mean of the observed VAF (left) and the clone-specific allele frequency *ϕ* (right) for each variant and clone. The ***π***-weighted VAF correlates with *ϕ*. **d**, Accessibility at the HIST1H cluster of replication-dependent histone genes in cells with low and high Clone 4 probability; several genes of the cluster ranked high for Clone 4. These genes are transcribed mainly during S phase(33). The adjusted p-value is descriptive (Methods).

**Fig. S5.**
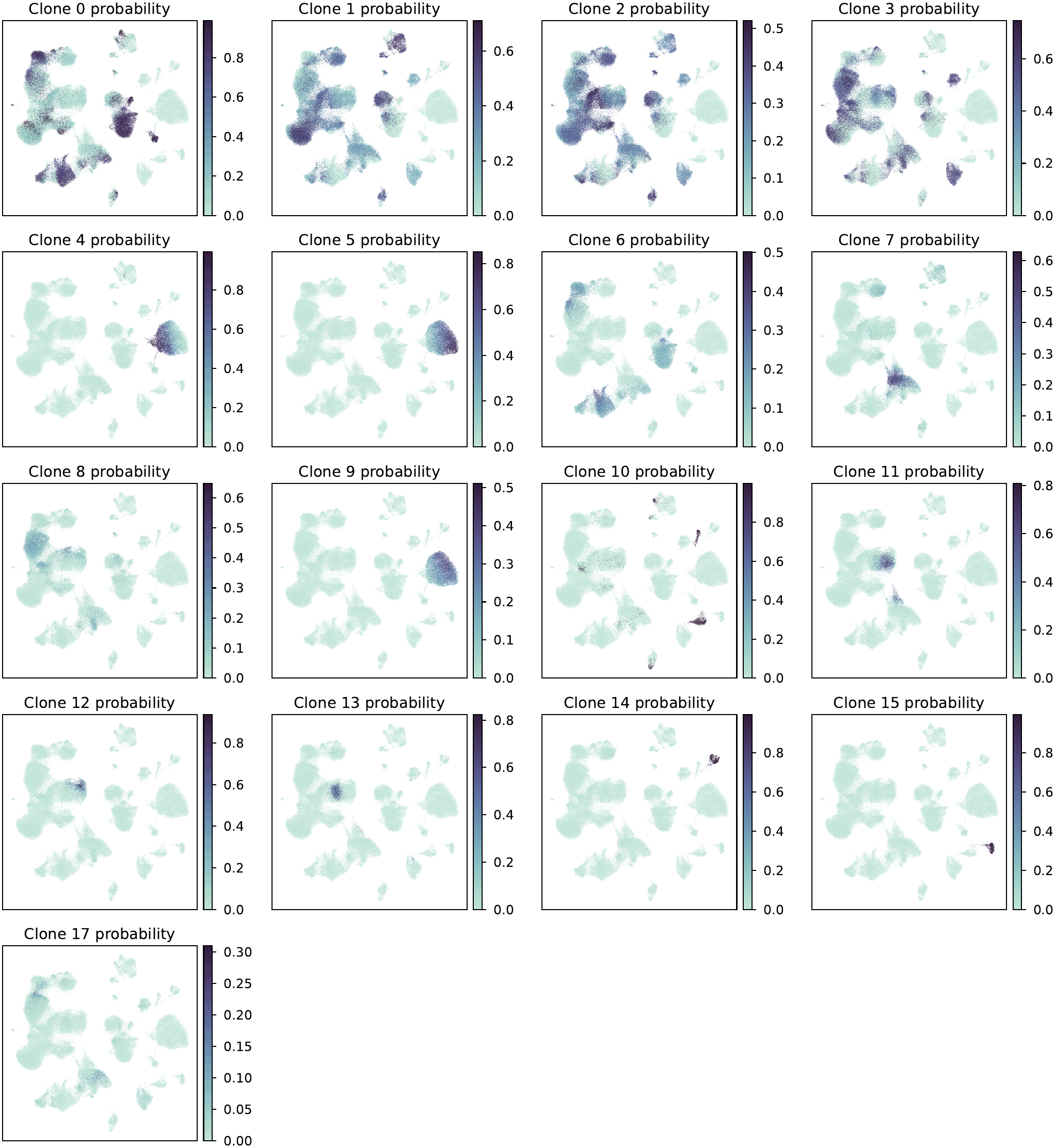
Clonal probabilities of the inferred clones in the ovarian cohort. The clonal probability ***π*** of each of the 17 inferred clones on the UMAP of the cell-state space, colored as in Fig. 5. The variants that each clone carries are given by its allele frequencies in Supplementary Fig. S7, and the observed VAF of these variants in Supplementary Fig. S6; several clones share a variant, for example Clones 4, 5 and 9 all carry 7738T>C, and Clones 5 and 9 additionally carry 12652T>C and 9554G>A, respectively. Clones 0 and 2 have no variant that exceeds the shared component by more than 0.02 and cover large parts of the state space.

**Fig. S6.**
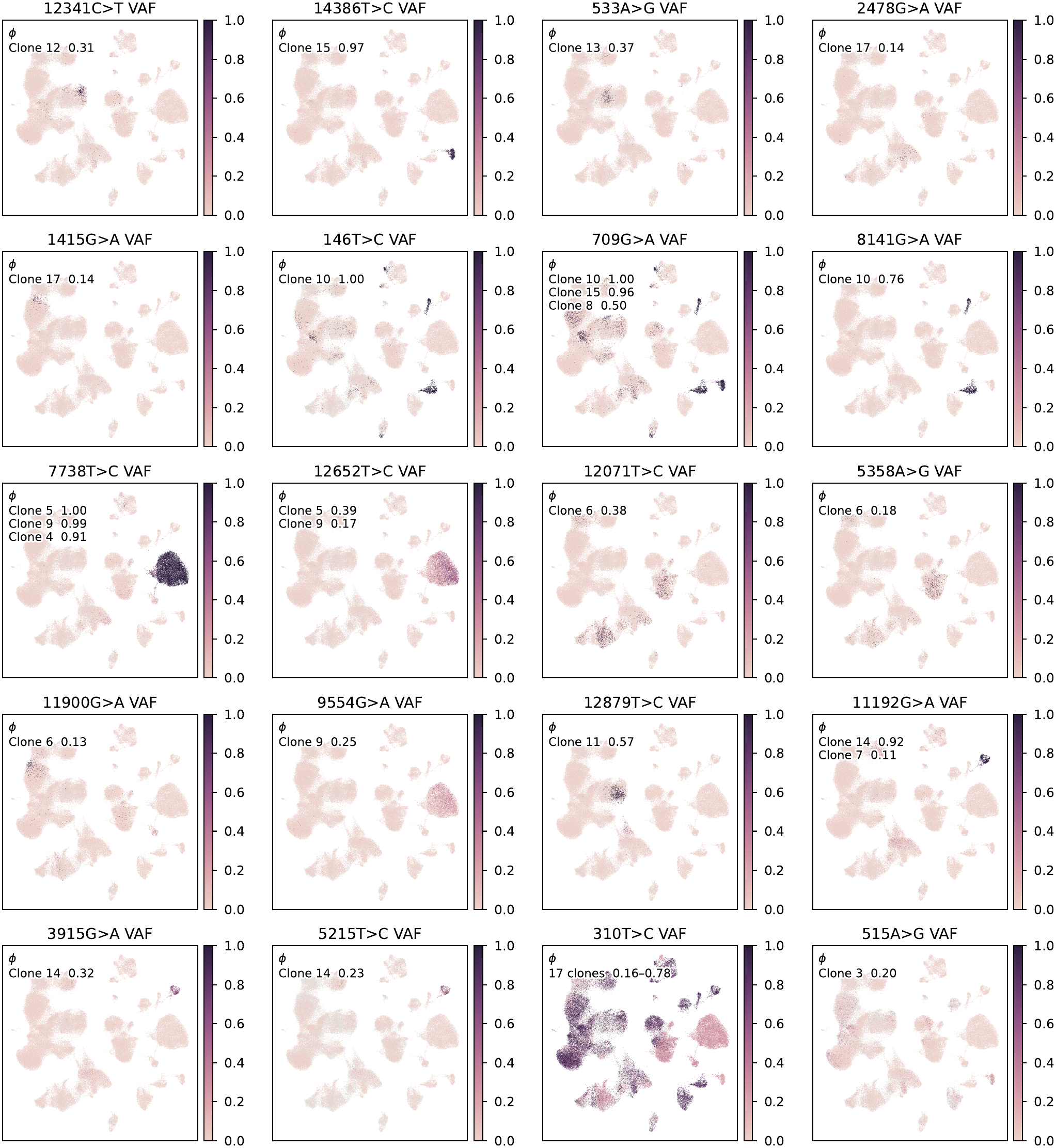
Observed VAF of the variants of Fig. 5d. The observed VAF of the 19 variants drawn in Fig. 5d, in the same order, and of 515A>G (not drawn there; Methods), on the UMAP of the cell-state space; gray, mtDNA depth below 10. Each panel lists the clones that carry the variant at *ϕ* ≥ 0.1 with their *ϕ*; 310T>C is carried by all 17 clones. The maps of 709G>A, 14386T>C, 7738T>C, 12652T>C and 9554G>A also appear in Supplementary Fig. S8, and that of 12879T>C in Fig. 5c, so that every variant behind Fig. 5d is shown. The color scale is 0–1 in every panel.

**Fig. S7.**
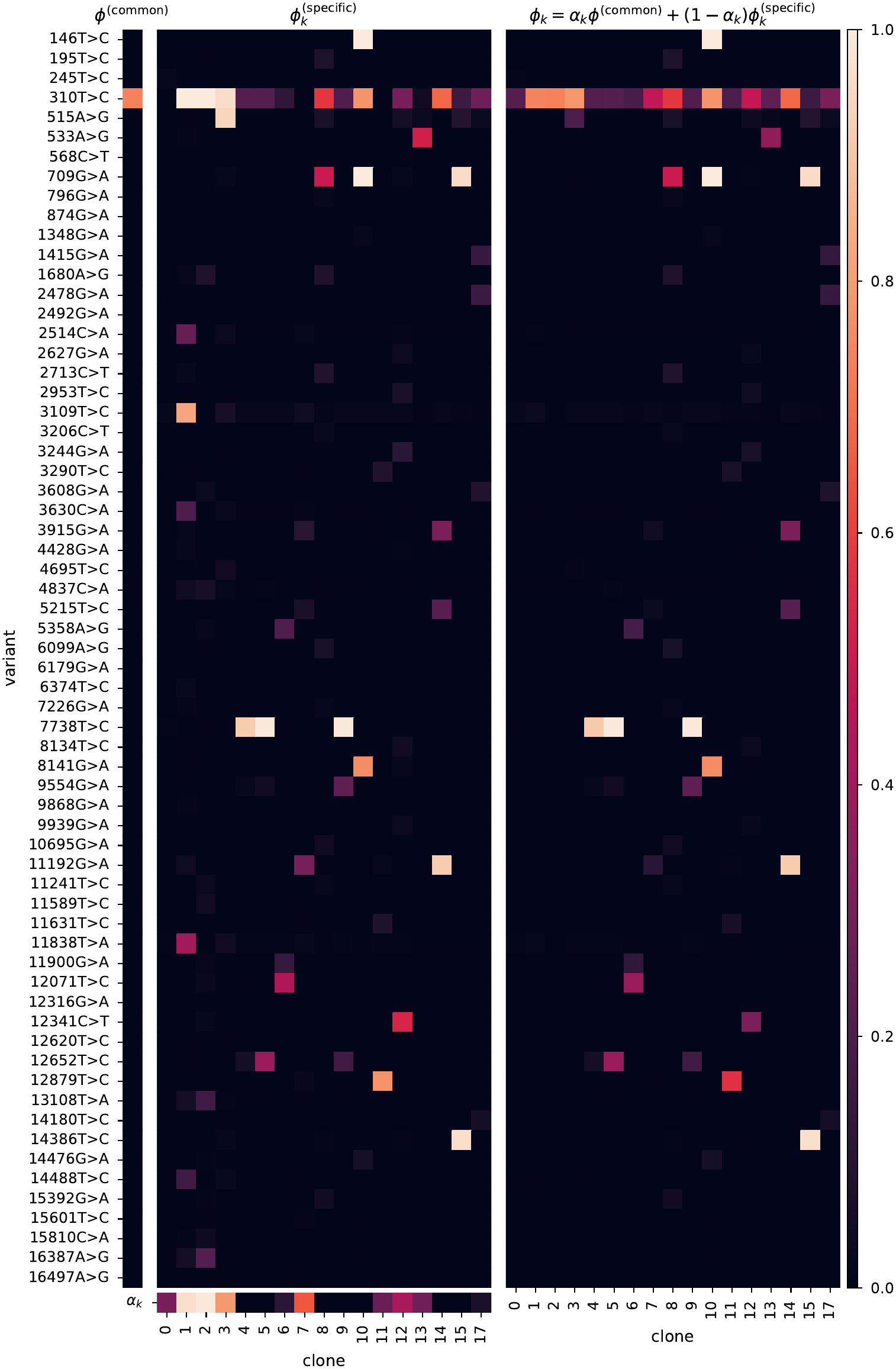
Decomposition of the allele frequency in the ovarian cohort. The shared component ***ϕ***^(common)^ (left; one value per variant), the clone-specific component 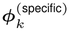 with the weight *α*_*k*_ of the mixture below it (middle; one value per clone), and the resulting allele frequency 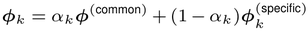 (right), for the 17 inferred clones and all 64 variants. The shared component is concentrated on 310T>C (0.74), which is carried by almost every covered cell, and 63 of the 64 variants are assigned a shared frequency below 0.01. Because the heteroplasmy of 310T>C differs between donors, the separation is partial rather than complete. Fitting the same data without this decomposition changed the relation between capture and cell-state enrichment only slightly (*ρ* = +0.83 and partial *ρ* = +0.71, against +0.81 and +0.75 with it); the two fits differ in the reassignment of the shared variant and in the refitting of the cell state, so this comparison shows that the conclusions do not depend on the decomposition rather than that the decomposition has no effect.

**Fig. S8.**
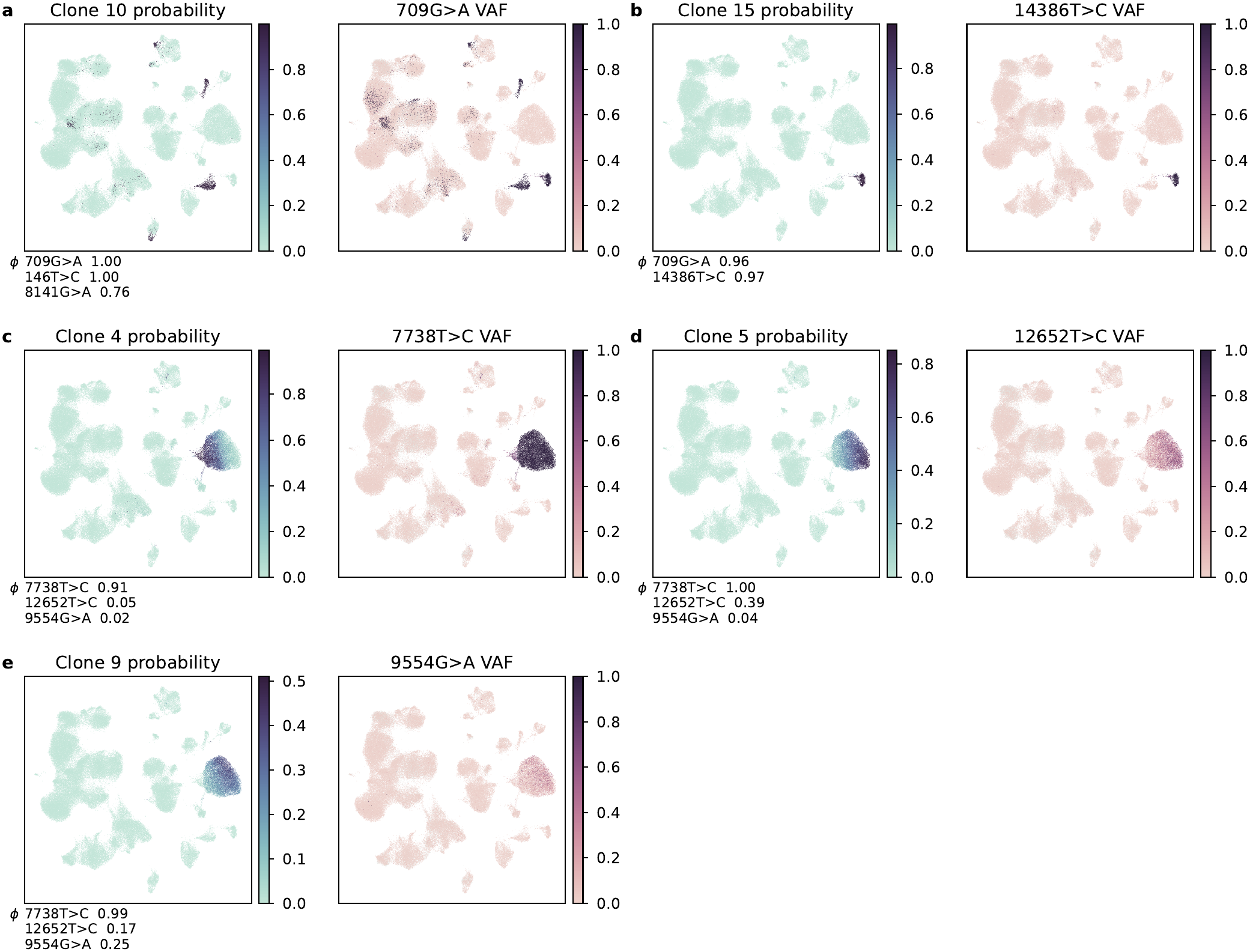
Clones that share a variant in the ovarian cohort. The clonal probability ***π*** of five inferred clones (as in Supplementary Fig. S5), each beside the observed VAF of a variant that it carries; the allele frequencies of the clone are given in each ***π*** panel, and the observed VAF of all variants of Fig. 5d in Supplementary Fig. S6. Most inferred clones of the cohort are confined to a single donor (10 of the 17 have more than 90% of their clonal probability in one donor), which is consistent with the donor being an explicit input to the decoder for ***π*. a**,**b**, One variant, 709G>A, carried by two clones in two donors: Clone 10 in SU-O-003, where the clone also carries 146T>C and 8141G>A, and Clone 15 in SU-O-001, which also carries 14386T>C, a variant of that donor alone. Each has essentially all of its clonal probability in the donor named. **c–e**, The lineage of donor SU-O-004: Clones 4, 5 and 9 all carry 7738T>C and differ in 12652T>C and in 9554G>A. Clones 5 and 9 occupy nearly the same region of the cell-state space (their ***π*** correlate at 0.91 within the donor) and are told apart by their allele frequencies rather than by their position, which is why Fig. 5d separates them and these maps do not. Color scales are per panel, because a variant observed at a mean VAF of 0.25 is not visible on a fixed 0–1 scale.

**Fig. S9.**
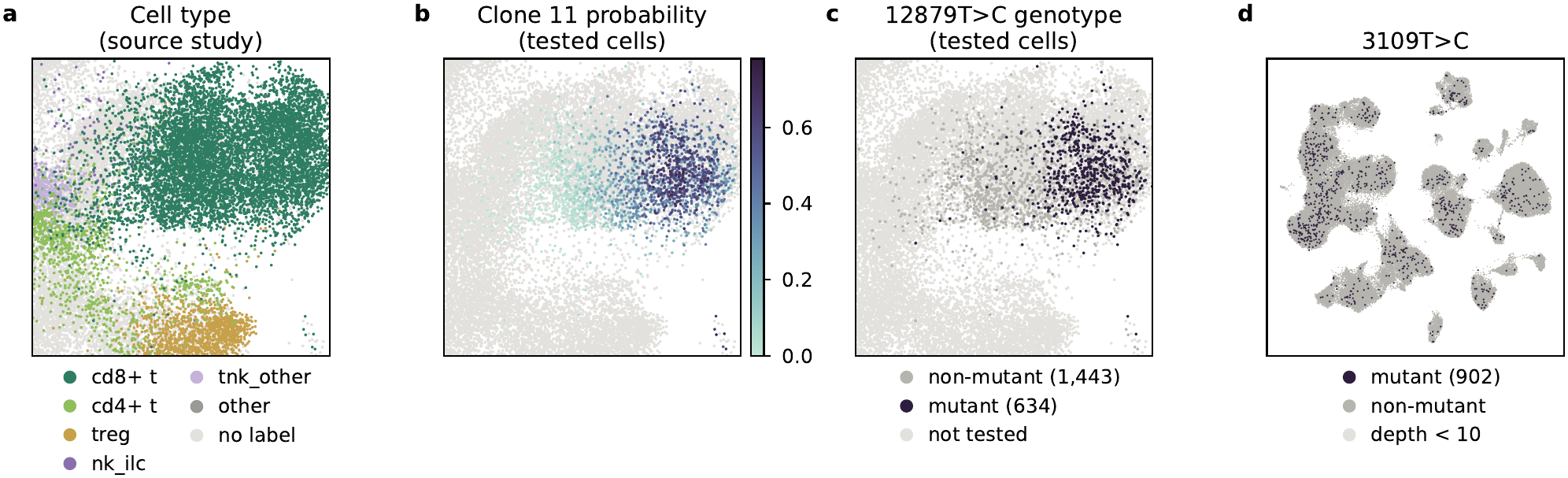
Cells entering the motif tests of Fig. 5f–h, and a variant without cell-state enrichment. **a–c**, The region of the cell-state space containing the tested cells, enlarged: cell type at the finer level of the source annotation (**a**), the clonal probability of Clone 11 in the tested cells, the 2,077 CD8^+^ T cells of one library of donor SU-O-005 covered at depth ≥ 10 (**b**), and the 12879T>C genotype of the same cells (**c**; mutant, VAF ≥ 0.10); other cells are drawn in light gray. **d**, The 902 mutant cells of 3109T>C on the cohort embedding, with covered non-mutant cells in gray, for comparison with the localized variant in Fig. 5c (right).

**Fig. S10.**
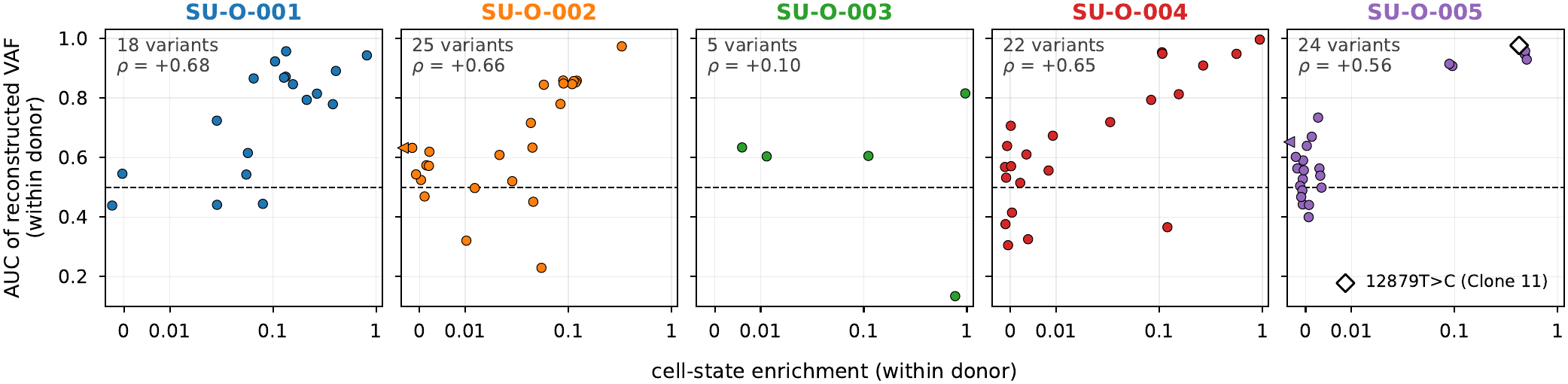
Capture and cell-state enrichment within each donor of the ovarian cohort. The AUC of the reconstructed VAF against cell-state enrichment, as in Fig. 5e (right), recomputed within each donor: the non-mutant cells are those of the same donor, and enrichment is computed on the covered cells of that donor (Methods). Each point is a variant with at least 15 mutant and 15 non-mutant cells in the donor, colored by donor as in Fig. 5a; *ρ*, Spearman correlation. SU-O-003 has only five variants (*ρ* = +0.10); among them, 1348G>A is enriched within the donor but not captured (AUC 0.13), although its AUC across the cohort is 0.89, an example of a variant that appears captured across the cohort only through the donor. The open diamond marks 12879T>C, the variant of Clone 11. 310T>C in SU-O-005, at an enrichment of −0.03, is drawn at the left edge.

**Supplementary Table 1.**
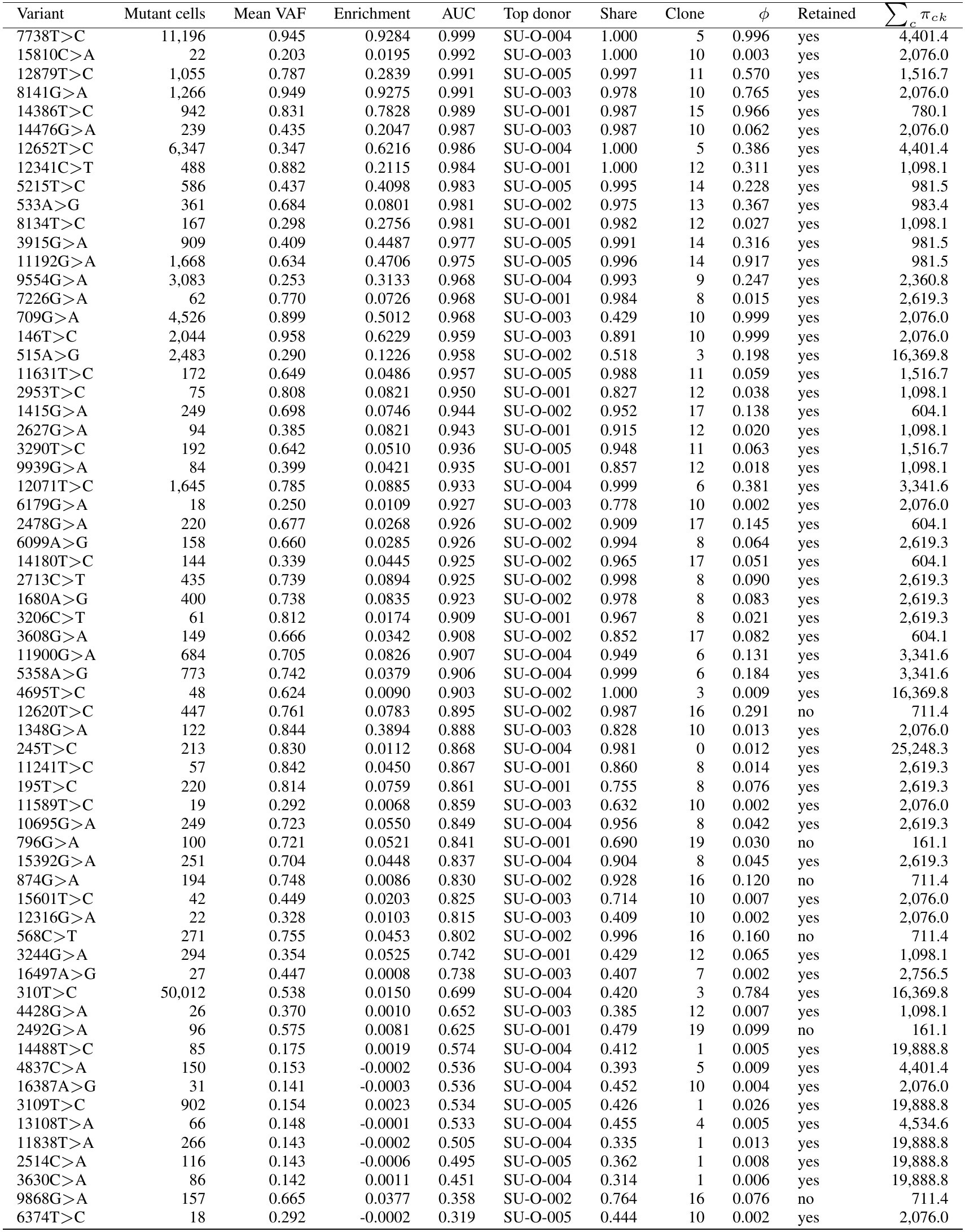
The 64 variants of the ovarian cohort (`SupplementaryTable_ovarian_variants.csv`): number of mutant cells, mean VAF of mutant cells, cell-state enrichment and AUC as in Fig. 5e, the donor carrying most mutant cells and its share of them, and the clone with the largest fitted allele frequency ***ϕ***, with that frequency, whether the clone was retained after pruning, and its total clonal probability ∑*c π*_*ck*_.

| Variant | Mutant cells | Mean VAF | Enrichment | AUC | Top donor | Share | Clone | ϕ | Retained | ∑<sub>c</sub> π<sub>ck</sub> |
|---|---|---|---|---|---|---|---|---|---|---|
| 7738T>C | 11,196 | 0.945 | 0.9284 | 0.999 | SU-O-004 | 1.000 | 5 | 0.996 | yes | 4,401.4 |
| 15810C>A | 22 | 0.203 | 0.0195 | 0.992 | SU-O-003 | 1.000 | 10 | 0.003 | yes | 2,076.0 |
| 12879T>C | 1,055 | 0.787 | 0.2839 | 0.991 | SU-O-005 | 0.997 | 11 | 0.570 | yes | 1,516.7 |
| 8141G>A | 1,266 | 0.949 | 0.9275 | 0.991 | SU-O-003 | 0.978 | 10 | 0.765 | yes | 2,076.0 |
| 14386T>C | 942 | 0.831 | 0.7828 | 0.989 | SU-O-001 | 0.987 | 15 | 0.966 | yes | 780.1 |
| 14476G>A | 239 | 0.435 | 0.2047 | 0.987 | SU-O-003 | 0.987 | 10 | 0.062 | yes | 2,076.0 |
| 12652T>C | 6,347 | 0.347 | 0.6216 | 0.986 | SU-O-004 | 1.000 | 5 | 0.386 | yes | 4,401.4 |
| 12341C>T | 488 | 0.882 | 0.2115 | 0.984 | SU-O-001 | 1.000 | 12 | 0.311 | yes | 1,098.1 |
| 5215T>C | 586 | 0.437 | 0.4098 | 0.983 | SU-O-005 | 0.995 | 14 | 0.228 | yes | 981.5 |
| 533A>G | 361 | 0.684 | 0.0801 | 0.981 | SU-O-002 | 0.975 | 13 | 0.367 | yes | 983.4 |
| 8134T>C | 167 | 0.298 | 0.2756 | 0.981 | SU-O-001 | 0.982 | 12 | 0.027 | yes | 1,098.1 |
| 3915G>A | 909 | 0.409 | 0.4487 | 0.977 | SU-O-005 | 0.991 | 14 | 0.316 | yes | 981.5 |
| 11192G>A | 1,668 | 0.634 | 0.4706 | 0.975 | SU-O-005 | 0.996 | 14 | 0.917 | yes | 981.5 |
| 9554G>A | 3,083 | 0.253 | 0.3133 | 0.968 | SU-O-004 | 0.993 | 9 | 0.247 | yes | 2,360.8 |
| 7226G>A | 62 | 0.770 | 0.0726 | 0.968 | SU-O-001 | 0.984 | 8 | 0.015 | yes | 2,619.3 |
| 709G>A | 4,526 | 0.899 | 0.5012 | 0.968 | SU-O-003 | 0.429 | 10 | 0.999 | yes | 2,076.0 |
| 146T>C | 2,044 | 0.958 | 0.6229 | 0.959 | SU-O-003 | 0.891 | 10 | 0.999 | yes | 2,076.0 |
| 515A>G | 2,483 | 0.290 | 0.1226 | 0.958 | SU-O-002 | 0.518 | 3 | 0.198 | yes | 16,369.8 |
| 11631T>C | 172 | 0.649 | 0.0486 | 0.957 | SU-O-005 | 0.988 | 11 | 0.059 | yes | 1,516.7 |
| 2953T>C | 75 | 0.808 | 0.0821 | 0.950 | SU-O-001 | 0.827 | 12 | 0.038 | yes | 1,098.1 |
| 1415G>A | 249 | 0.698 | 0.0746 | 0.944 | SU-O-002 | 0.952 | 17 | 0.138 | yes | 604.1 |
| 2627G>A | 94 | 0.385 | 0.0821 | 0.943 | SU-O-001 | 0.915 | 12 | 0.020 | yes | 1,098.1 |
| 3290T>C | 192 | 0.642 | 0.0510 | 0.936 | SU-O-005 | 0.948 | 11 | 0.063 | yes | 1,516.7 |
| 9939G>A | 84 | 0.399 | 0.0421 | 0.935 | SU-O-001 | 0.857 | 12 | 0.018 | yes | 1,098.1 |
| 12071T>C | 1,645 | 0.785 | 0.0885 | 0.933 | SU-O-004 | 0.999 | 6 | 0.381 | yes | 3,341.6 |
| 6179G>A | 18 | 0.250 | 0.0109 | 0.927 | SU-O-003 | 0.778 | 10 | 0.002 | yes | 2,076.0 |
| 2478G>A | 220 | 0.677 | 0.0268 | 0.926 | SU-O-002 | 0.909 | 17 | 0.145 | yes | 604.1 |
| 6099A>G | 158 | 0.660 | 0.0285 | 0.926 | SU-O-002 | 0.994 | 8 | 0.064 | yes | 2,619.3 |
| 14180T>C | 144 | 0.339 | 0.0445 | 0.925 | SU-O-002 | 0.965 | 17 | 0.051 | yes | 604.1 |
| 2713C>T | 435 | 0.739 | 0.0894 | 0.925 | SU-O-002 | 0.998 | 8 | 0.090 | yes | 2,619.3 |
| 1680A>G | 400 | 0.738 | 0.0835 | 0.923 | SU-O-002 | 0.978 | 8 | 0.083 | yes | 2,619.3 |
| 3206C>T | 61 | 0.812 | 0.0174 | 0.909 | SU-O-001 | 0.967 | 8 | 0.021 | yes | 2,619.3 |
| 3608G>A | 149 | 0.666 | 0.0342 | 0.908 | SU-O-002 | 0.852 | 17 | 0.082 | yes | 604.1 |
| 11900G>A | 684 | 0.705 | 0.0826 | 0.907 | SU-O-004 | 0.949 | 6 | 0.131 | yes | 3,341.6 |
| 5358A>G | 773 | 0.742 | 0.0379 | 0.906 | SU-O-004 | 0.999 | 6 | 0.184 | yes | 3,341.6 |
| 4695T>C | 48 | 0.624 | 0.0090 | 0.903 | SU-O-002 | 1.000 | 3 | 0.009 | yes | 16,369.8 |
| 12620T>C | 447 | 0.761 | 0.0783 | 0.895 | SU-O-002 | 0.987 | 16 | 0.291 | no | 711.4 |
| 1348G>A | 122 | 0.844 | 0.3894 | 0.888 | SU-O-003 | 0.828 | 10 | 0.013 | yes | 2,076.0 |
| 245T>C | 213 | 0.830 | 0.0112 | 0.868 | SU-O-004 | 0.981 | 0 | 0.012 | yes | 25,248.3 |
| 11241T>C | 57 | 0.842 | 0.0450 | 0.867 | SU-O-001 | 0.860 | 8 | 0.014 | yes | 2,619.3 |
| 195T>C | 220 | 0.814 | 0.0759 | 0.861 | SU-O-001 | 0.755 | 8 | 0.076 | yes | 2,619.3 |
| 11589T>C | 19 | 0.292 | 0.0068 | 0.859 | SU-O-003 | 0.632 | 10 | 0.002 | yes | 2,076.0 |
| 10695G>A | 249 | 0.723 | 0.0550 | 0.849 | SU-O-004 | 0.956 | 8 | 0.042 | yes | 2,619.3 |
| 796G>A | 100 | 0.721 | 0.0521 | 0.841 | SU-O-001 | 0.690 | 19 | 0.030 | no | 161.1 |
| 15392G>A | 251 | 0.704 | 0.0448 | 0.837 | SU-O-004 | 0.904 | 8 | 0.045 | yes | 2,619.3 |
| 874G>A | 194 | 0.748 | 0.0086 | 0.830 | SU-O-002 | 0.928 | 16 | 0.120 | no | 711.4 |
| 15601T>C | 42 | 0.449 | 0.0203 | 0.825 | SU-O-003 | 0.714 | 10 | 0.007 | yes | 2,076.0 |
| 12316G>A | 22 | 0.328 | 0.0103 | 0.815 | SU-O-003 | 0.409 | 10 | 0.002 | yes | 2,076.0 |
| 568C>T | 271 | 0.755 | 0.0453 | 0.802 | SU-O-002 | 0.996 | 16 | 0.160 | no | 711.4 |
| 3244G>A | 294 | 0.354 | 0.0525 | 0.742 | SU-O-001 | 0.429 | 12 | 0.065 | yes | 1,098.1 |
| 16497A>G | 27 | 0.447 | 0.0008 | 0.738 | SU-O-003 | 0.407 | 7 | 0.002 | yes | 2,756.5 |
| 310T>C | 50,012 | 0.538 | 0.0150 | 0.699 | SU-O-004 | 0.420 | 3 | 0.784 | yes | 16,369.8 |
| 4428G>A | 26 | 0.370 | 0.0010 | 0.652 | SU-O-003 | 0.385 | 12 | 0.007 | yes | 1,098.1 |
| 2492G>A | 96 | 0.575 | 0.0081 | 0.625 | SU-O-001 | 0.479 | 19 | 0.099 | no | 161.1 |
| 14488T>C | 85 | 0.175 | 0.0019 | 0.574 | SU-O-004 | 0.412 | 1 | 0.005 | yes | 19,888.8 |
| 4837C>A | 150 | 0.153 | -0.0002 | 0.536 | SU-O-004 | 0.393 | 5 | 0.009 | yes | 4,401.4 |
| 16387A>G | 31 | 0.141 | -0.0003 | 0.536 | SU-O-004 | 0.452 | 10 | 0.004 | yes | 2,076.0 |
| 3109T>C | 902 | 0.154 | 0.0023 | 0.534 | SU-O-005 | 0.426 | 1 | 0.026 | yes | 19,888.8 |
| 13108T>A | 66 | 0.148 | -0.0001 | 0.533 | SU-O-004 | 0.455 | 4 | 0.005 | yes | 4,534.6 |
| 11838T>A | 266 | 0.143 | -0.0002 | 0.505 | SU-O-004 | 0.335 | 1 | 0.013 | yes | 19,888.8 |
| 2514C>A | 116 | 0.143 | -0.0006 | 0.495 | SU-O-005 | 0.362 | 1 | 0.008 | yes | 19,888.8 |
| 3630C>A | 86 | 0.142 | 0.0011 | 0.451 | SU-O-004 | 0.314 | 1 | 0.006 | yes | 19,888.8 |
| 9868G>A | 157 | 0.665 | 0.0377 | 0.358 | SU-O-002 | 0.764 | 16 | 0.076 | no | 711.4 |
| 6374T>C | 18 | 0.292 | -0.0002 | 0.319 | SU-O-005 | 0.444 | 10 | 0.002 | yes | 2,076.0 |

**Supplementary Table 2.** Settings used to fit CloneVI in each analysis. Other training settings, such as the learning rate, the number of epochs and the random seed, are provided with the code.

|  | Simulation | MAESTER | CLL | Ovarian |
|---|---|---|---|---|
| Variants (*M*) | 6 | 26 | 18 | 64 |
| Candidate clones (*K*) | 10 | 20 | 20 | 20 |
| Sample indicator | none | none | none | donor (5) |
| Decomposition of ***ϕ*** | no | no | yes | yes |
| Regularization (*γ*<sub>*π*</sub>, *γ*<sub>***ϕ***</sub>) | none | none | 0.01, 1 | 0.01, 1 |

**Supplementary Table 3.** Simulation settings (Fig. 2).

|  |  |
|---|---|
| Cells and expression | Scanpy PBMC3k (processed): 2,638 cells, 1,838 genes (raw counts), eight Louvain clusters |
| Clones | 2 |
| Loci (*M*) | 6 |
| Mutation profiles | **r**<sub>0</sub> = (0, 1, 0, 0, 1, 1), **r**<sub>1</sub> = (0, 1, 1, 1, 0, 0) |
| Depth | *d*<sub>*cm*</sub> ∼ Poisson(80) |
| Mutation counts | **b**<sub>*c*</sub> = **r**<sub>*ρ*<sub>*c*</sub></sub> ⊙ **d**<sub>*c*</sub> (no background, no dropout) |
| Clonal probability per cluster | fixed: CD4 T cells (1.00, 0.00); CD14<sup>+</sup> monocytes (0.00, 1.00); B cells (0.50, 0.50);<br>drawn: CD8 T cells (0.43, 0.57); NK cells (0.79, 0.21); FCGR3A<sup>+</sup> monocytes (0.42, 0.58); dendritic cells (0.50, 0.50); megakaryocytes (0.51, 0.49) |
| Resulting clones | Clone 0: 1,676 cells; Clone 1: 962 cells |
| Random seeds | drawn cluster probabilities: 1234; clone assignment and read counts: not seeded<br>(the generated data are provided) |

**Supplementary Table 4.**
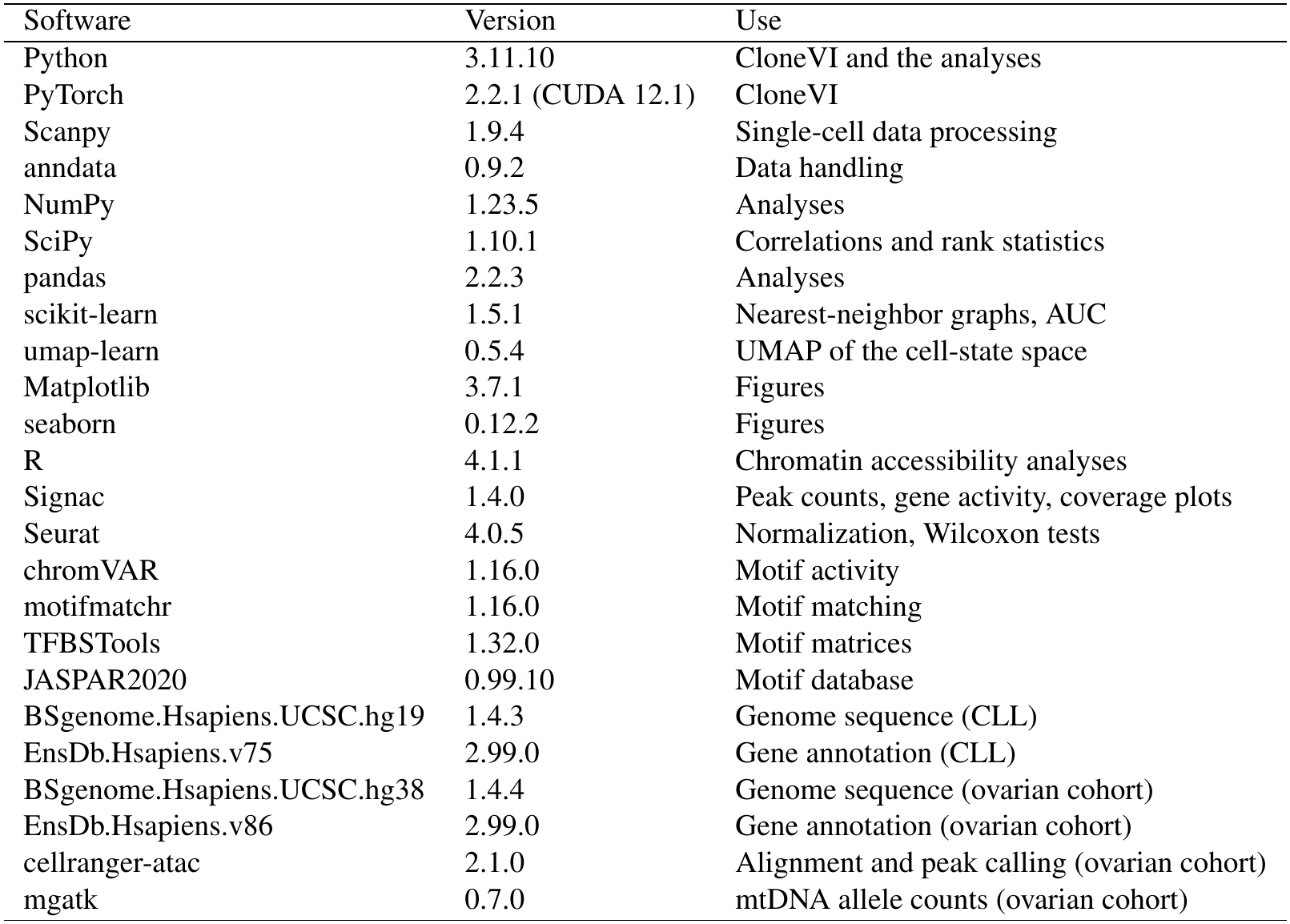
Software used in this study. The versions of all packages are listed with the code.

| Software | Version | Use |
|---|---|---|
| Python | 3.11.10 | CloneVI and the analyses |
| PyTorch | 2.2.1 (CUDA 12.1) | CloneVI |
| Scanpy | 1.9.4 | Single-cell data processing |
| anndata | 0.9.2 | Data handling |
| NumPy | 1.23.5 | Analyses |
| SciPy | 1.10.1 | Correlations and rank statistics |
| pandas | 2.2.3 | Analyses |
| scikit-learn | 1.5.1 | Nearest-neighbor graphs, AUC |
| umap-learn | 0.5.4 | UMAP of the cell-state space |
| Matplotlib | 3.7.1 | Figures |
| seaborn | 0.12.2 | Figures |
| R | 4.1.1 | Chromatin accessibility analyses |
| Signac | 1.4.0 | Peak counts, gene activity, coverage plots |
| Seurat | 4.0.5 | Normalization, Wilcoxon tests |
| chromVAR | 1.16.0 | Motif activity |
| motifmatchr | 1.16.0 | Motif matching |
| TFBSTools | 1.32.0 | Motif matrices |
| JASPAR2020 | 0.99.10 | Motif database |
| BSgenome.Hsapiens.UCSC.hg19 | 1.4.3 | Genome sequence (CLL) |
| EnsDb.Hsapiens.v75 | 2.99.0 | Gene annotation (CLL) |
| BSgenome.Hsapiens.UCSC.hg38 | 1.4.4 | Genome sequence (ovarian cohort) |
| EnsDb.Hsapiens.v86 | 2.99.0 | Gene annotation (ovarian cohort) |
| cellranger-atac | 2.1.0 | Alignment and peak calling (ovarian cohort) |
| mgatk | 0.7.0 | mtDNA allele counts (ovarian cohort) |

